# SPOP targets the immune transcription factor IRF1 for proteasomal degradation

**DOI:** 10.1101/2022.10.10.511567

**Authors:** Milica Vunjak, Irene Schwartz, Adriana Cantoran García, Marialaura Mastrovito, Matthias Hinterndorfer, Melanie de Almeida, Valentina Budroni, Jingkui Wang, Kimon Froussios, Julian Jude, Thomas Decker, Johannes Zuber, Gijs A. Versteeg

## Abstract

Adaptation of the functional proteome is essential to counter pathogens during infection, yet precisely timed degradation of these response proteins after pathogen clearance is likewise key to preventing autoimmunity. Interferon Regulatory Factor 1 (IRF1) plays an essential role as a transcription factor in driving the expression of immune response genes during infection. The striking difference in functional output with other IRFs, is that IRF1 also drives the expression of various cell cycle inhibiting factors, making it an important tumor suppressor. Thus, it is critical to regulate the abundance of IRF1 to achieve a “Goldilocks” zone in which there is sufficient IRF1 to prevent tumorigenesis, yet not too much which could drive excessive immune activation. Using genetic screening, we identified the E3 ligase receptor Speckle Type BTB/POZ Protein (SPOP) to mediate IRF1 proteasomal turnover. We identified four S/T-rich degrons in IRF1 required for its SPOP MATH domain-dependent turnover. In the absence of SPOP, increased IRF1 protein levels functionally drive increased transcription of IRF1-response-genes, underpinning the biological significance of SPOP in curtailing IRF1 protein abundance.

## Introduction

Adaptation of the functional proteome is essential to counter pathogens during infection, yet precisely timed degradation of these response proteins after pathogen clearance is likewise key to preventing autoimmunity and hyperinflammation (Crow & Stetson, 2022).

Recognition of Pathogen Associated Molecular Patterns (PAMPs), and subsequent secretion of cytokines known as interferons (IFN), are particularly critical for conferring an anti-pathogen response (Takeuchi & Akira, 2010). Type I interferons (IFNα/ß) can be secreted by most cells upon infection, whereas type II IFNs (IFNγ) are secreted by specific cell types such as Natural Killer (NK) cells, and T helper 1 (T_H_1) cells. Binding of these cytokines to their receptors initiates cell signaling driving transcription of hundreds of response genes, which collectively mediate the anti-pathogen response (Schoggins, 2019).

Interferon Regulatory Factors (IRFs) play essential roles as transcription factors in the IFN system. Some IRFs, such as IRF3, are constitutively expressed, their activity is predominantly regulated through phosphorylation, and they are essential for induction of type I IFNs (Tamura et al., 2008). In contrast, some other IRFs, including IRF1, are transcriptionally induced during infection (Feng et al., 2021).

IRF1 is a key family member for the induction of the innate immune response (Forero et al., 2019; Kimura et al., 1994; Langlais et al., 2016). The importance to precisely control IRF1 levels and its functional output is underpinned by the fact that aberrant IRF1 expression is linked to inflammatory syndromes, such as Behcet’s disease, arthritis and systemic lupus erythematosus (Donn et al., 2001; Govind et al., 2014; Lee et al., 2007; Uddin et al., 2011; Z. Zhang et al., 2015). The striking difference in functional output with other IRFs is that IRF1 also drives the expression of various cell cycle inhibiting factors (e.g. p21) (Armstrong et al., 2012; Tanaka et al., 1996), making it an important tumor suppressor (Nozawa et al., 1999; Xie et al., 2003). Thus, it is critical to regulate the abundance of IRF1 to achieve “Goldilocks” levels at which there is sufficient IRF1 to prevent tumorigenesis, yet not too much which could cause excessive immune activation.

In addition to its transcriptional regulation, the other main means of IRF1 regulation is at a protein level through rapid degradation by the proteasome (Nakagawa & Yokosawa, 2000; Pion et al., 2009), which seems to in turn be critical to limit transcriptional output and reduce the IRF1 protein pools once IRF1 transcription is shut off during infection resolution. In line with this notion, proteasome inhibitors have been shown to block IRF1 degradation (Attard et al., 2005). Several factors have thus far been reported to mediate poly-ubiquitination of IRF1, and direct its proteasomal degradation.

The importance of these individual regulators for IRF1 turnover is to a degree determined by their cellular context (Liu et al., 2021; Remoli et al., 2020). This draws an interesting parallel to other transcription factors which can be ubiquitinated by various different E3 ligase complexes, depending on cellular context. cMYC is a prime example of such a cell-context regulated transcription factor. A Cullin E3 ligase complex using the FBXW7 substrate receptor module (SCF^FBXW7^) has been reported as a key factor in cMYC degradative ubiquitination in many cell types (Welcker et al., 2004; Yada et al., 2004). In contrast, various others, including TRIM32, CHIP, and several other Cullin F-box complexes, have been shown to be important for cMYC degradation, yet in more cell-specific contexts (Sun et al., 2021). Identification and characterization of such E3 ligases is critical for ultimately understanding how their action controls transcriptional output in various cell types, within the context of whole organisms.

As for other transcription factors, turnover mechanisms can be different for distinct cellular IRF1 protein pools, dependent on their localization and functional engagement in transcription. As such, understanding which functional IRF1 domains determine its turnover is important. IRF1 has an N-terminal DNA binding domain, followed by a nuclear localization signal (NLS), IRF1 transactivation IRF-association domain (IAD2), and a C-terminal intrinsically disordered enhancer region (IDR). This C-terminal IDR is an important determinant of IRF1 turnover (Nakagawa & Yokosawa, 2000). On the one hand, the IDR is stabilized by the chaperones HSP70 and HSP90 (Landré et al., 2013; Nakagawa & Yokosawa, 2000; Narayan et al., 2009, 2015, p. 7; Narayan, Halada, et al., 2011; Narayan, Pion, et al., 2011), whereas it is targeted for degrative ubiquitination by the protein quality control E3 ligase CHIP (Narayan et al., 2015; Narayan, Pion, et al., 2011). This has suggested that cells may, at least in part, turn over IRF1 co- or post-translationally using their general protein quality control pathways.

In addition, the N-terminal DNA binding domain has been reported to determine IRF1 stability (Landré et al., 2013). Consistent with a model in which IRF1 is ubiquitinated by the E3 ligase MDM2 in an unbound state, DNA binding competes with E3 ligase interaction, thereby stabilizing IRF1 in a functionally engaged state (Landré et al., 2013).

Lastly, the IAD2 domain has been implicated in driving IRF1 ubiquitination and degradation (Garvin et al., 2019; Pion et al., 2009). In part, this has been attributed to generation of a GSK3β-dependent phospho-degron on DNA-bound IRF1, and subsequent recognition by the SCF^FBXW7^ E3 ligase complex driving its degradation.

In this study, we set out to identify novel factors that control IRF1 abundance in an unbiased manner. Genome-wide genetic screens were performed to identify novel IRF1-specific E3 ligases. This identified the Cullin ubiquitin E3 ligase receptor SPOP to functionally mediate IRF1 degradation. Follow-up analyses identified four degrons in, or near the IRF1 IAD2 enhancer domain responsible for SPOP binding, and subsequent IRF1 ubiquitination and degradation. Consequently, loss of *SPOP* increased transcriptional output of IRF1-stimulated genes.

## Results

### Establishment of an IRF1 protein abundance reporter cell line for genetic screening

The IRF1 protein is ubiquitinated and turned over by the proteasome. However, the cellular E3 ligases contributing to targeting IRF1 for degradation have remained poorly understood. The overarching goal of this study was to identify and characterize novel regulators of IRF1 protein abundance. For the identification of IRF1 stability regulators, we characterized IRF1 degradation properties in the human RKO colon carcinoma cell line, as it is highly amenable for genetic screening.

IRF1 is transcriptionally induced upon type I or type II IFN stimulation in a wide variety of cells. Likewise, stimulation of RKO cells with IFNγ strongly induced endogenous IRF1 protein levels (Fig. 1A; 11.2-fold). Subsequent treatment with the translation inhibitor cycloheximide (CHX) rapidly chased out the induced IRF1 protein (Fig. 1A). In the absence of *de novo* protein synthesis, IRF1 levels returned to pre-stimulation protein levels within 4 h, indicating that it is highly unstable. In line with this finding, endogenous IRF1 protein levels measured by intracellular staining were also induced by IFNγ and depleted upon translation inhibition (Fig. 1B; left panel), whereas inhibition of proteasomal degradation by MG132 increased IRF1 levels (Fig. 1B; right panel). Together, these results indicate that IRF1 is induced by IFNγ and degraded by the proteasome in RKO cells, as has been described for other cell types, making it a suitable model for identification of IRF1 stability regulators.

**Figure 1.**
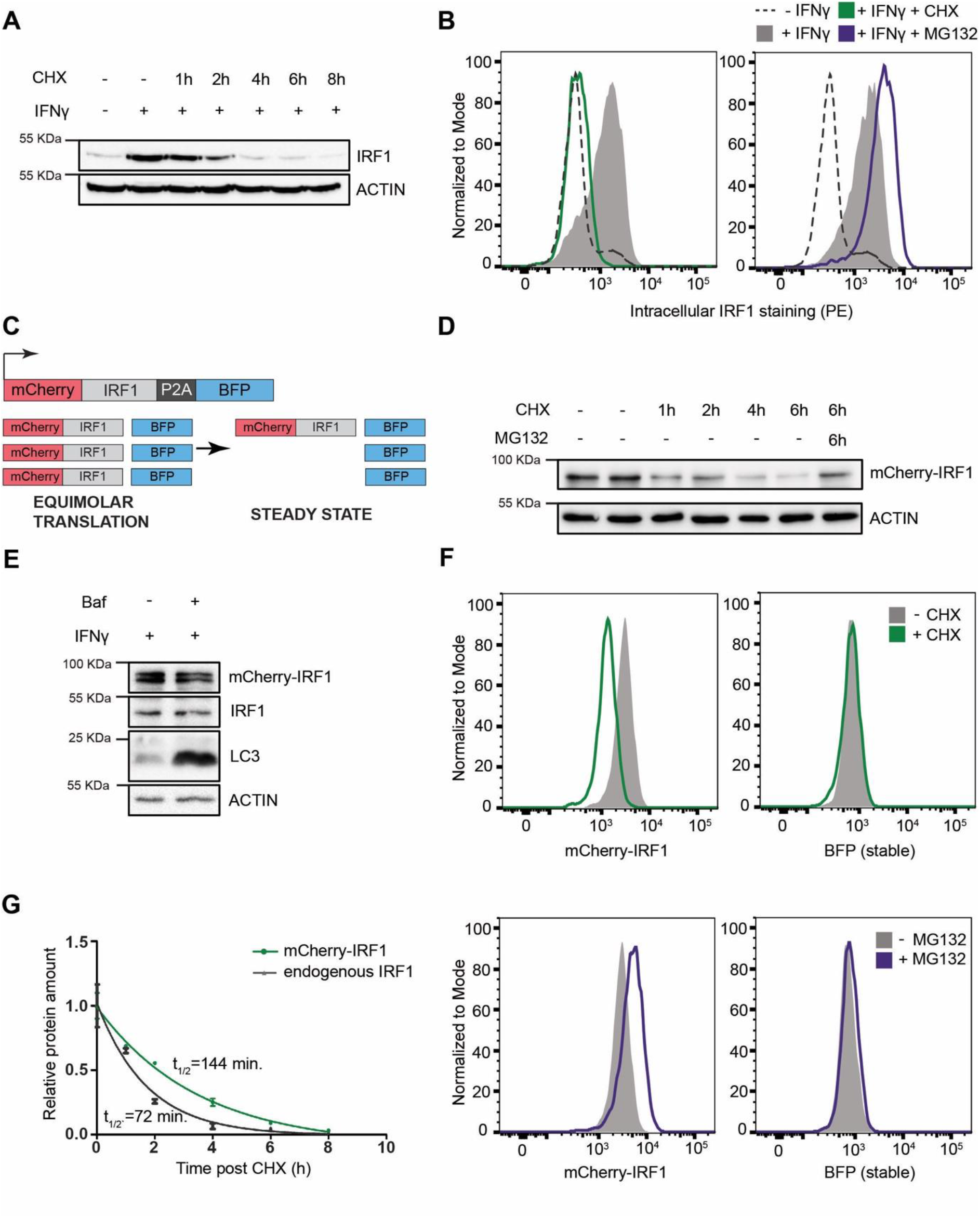
Establishment of an IRF1 protein abundance reporter cell line for genetic screening. RKO cells were stimulated with IFNγ for 4 h., and subsequently incubated with the translation inhibitor cycloheximide (CHX) in the absence or presence of proteasome inhibitor MG132, after which endogenous IRF1 levels were analyzed by **A)** Western blot, or **B)** intracellular staining and flow cytometry. **C)** A dual-color reporter construct, from which unstable mCherry-IRF1 and stable BFP are synthesized in equimolar amounts, yet mCherry-IRF1 accumulates at low steady-state protein concentrations resulting from it proteasomal degradation. **D)** RKO-mCherry-IRF1-P2A-BFP cells were incubated with CHX for the indicated times in the absence or presence of proteasome inhibitor MG132, after which mCherry-IRF1 levels were analyzed by Western blot through detection of IRF1 protein. **E)** RKO-mCherry-IRF1-P2A-BFP cells were stimulated with IFNγ for 4 h., and subsequently incubated with lysosomal degradation inhibitor Bafilomycin A (Baf) for 4 h., after which IRF1 levels were determined by Western blot. **F)** RKO-mCherry-IRF1-P2A-BFP cells were incubated with CHX or MG132 for 4 h., after which mCherry-IRF and BFP protein levels were analyzed by flow cytometry. **G)** RKO-mCherry-IRF1-P2A-BFP cells were incubated with CHX for the indicated times. Non-saturated Western blot signals for mCherry-IRF1 and endogenous IRF1 protein were quantified, normalized to ACTIN levels, plotted, and used for half-life calculation. n = 2.

Endogenous IRF1 protein levels are influenced by transcriptional and post-translational effects. Therefore, a strategy was devised to uncouple IRF1 protein abundance from its endogenous transcriptional program, thereby aiming to maximize the identification of regulators of protein degradation. To this end, a monoclonal cell line harboring Dox-inducible Cas9 was established, and a dual-color reporter construct under the control of a constitutively active promoter (Fig. 1C), consisting of an mCherry-IRF1 cassette separated from BFP by a P2A ribosomal skip site. Although both proteins are translated from a single transcript in equimolar quantities, the unstable mCherry-IRF1 was predicted to accumulate at lower steady-state levels than the stable BFP functioning as an internal control.

We reasoned that it would be important to screen in parallel with an unrelated protein that is degraded in a proteasome-dependent manner, in order to distinguish between general players in the ubiquitin proteasome system and factors specifically regulating IRF1 abundance. cMYC is constantly turned over by the proteasome with a 20-30 min. half-life (Gregory & Hann, 2000; Ramsay et al., 1986). Therefore, a comparable cell line expressing mCherry-cMYC was established and characterized alongside the mCherry-IRF1 cell line.

To investigate whether the mCherry-tagged IRF1 reporter had comparable degradation characteristics as its endogenous counterpart, cells were subjected to a CHX chase to determine protein stability. In line with endogenous IRF1, mCherry-IRF1 levels rapidly diminished within four hours (Fig. 1D). IRF1 degradation was prevented by simultaneous inhibition of the proteasome (Fig. 1D), whereas inhibition of autophagy with Bafilomycin A (Baf) did neither stabilize endogenous IRF1, nor mCherry-IRF1 (Fig. 1E). Similarly, mCherry-cMYC and endogenous cMYC were rapidly lost upon CHX treatment, which was prevented by simultaneous proteasome inhibition (Fig. S1A). This indicated that mCherry-IRF1, mCherry-cMYC, and their endogenous equivalents are all predominantly degraded by the proteasome.

Next, we set out to determine whether our fluorescent reporters were compatible with fluorescence-activated cell sorting (FACS)-based genetic screening. In particular, we assessed by flow cytometry whether inhibition of protein synthesis or proteasomal degradation would specifically affect the mCherry-tagged unstable proteins, without affecting their stable BFP internal controls. Consistent with our findings by Western blot readout, flow cytometry analysis also showed that both mCherry-IRF1 and mCherry-cMYC were degraded in the absence of *de novo* protein synthesis (Fig. 1F, Fig. S1B; top panels), and stabilized by proteasome inhibition (Fig. 1F, Fig. S1B; bottom panels), without affecting levels of the stable BFP internal control.

To determine IRF1 and cMYC half-lives, we quantified mCherry-IRF1 and IFNγ-induced endogenous IRF1 levels over time following inhibition of protein synthesis. Single exponential decay curve fitting indicated the half-life of endogenous IRF1 to be 72 min., whereas the half-life of mCherry-IRF1 was twice as long at 144 min. (Fig. 1G). Importantly, IFNγ treatment did not alter constitutively expressed mCherry-IRF1 levels (Fig. S1C), indicating that its degradation machinery is likely already active under non-stimulated conditions. Similar measurements for mCherry-cMYC indicated a 24 min. half-life for mCherry-cMYC and 21 min. for endogenous cMYC in the same cells (Fig. S1D), consistent with previously published work (Welcker et al., 2004; Yada et al., 2004). These combined data indicate that the N-terminal mCherry fusions of IRF1 and cMYC phenocopied the proteasomal degradation and instability of their endogenous counterparts.

Lastly, we assessed whether our selected monoclonal mCherry-IRF1 and mCherry-cMYC reporter cell lines had retained the ability to drive genome editing. To this end, cells were transduced with a lentiviral vector constitutively expressing an sgRNA targeting the cell-essential factor Ribonucleotide Reductase Catalytic M1 (*RRM1*), while co-expressing iRFP. Subsequently, the ratio between these transduced cells and non-transduced cells in the same cell pool were measured over time by flow cytometry as a proxy for cell fitness.

As anticipated, the presence of Dox in the culture medium to induce Cas9 resulted in a near-complete loss of iRFP^+^ sg*RRM1* cells from the cell pool in six days in both cell lines (Fig. S1E/F), whereas parallel sg*AAVS1* control samples remained at a constant ratio. In contrast, in the absence of Dox, sg*RRM1* cells were not depleted over the two-week course of the experiment. This indicated that the mCherry-IRF1 and mCherry-cMYC cell lines had minimal editing in the absence of Dox, yet efficient genome editing upon Dox-induced Cas9 expression.

Taken together, our data demonstrate that mCherry-IRF1 and mCherry-cMYC phenocopy the unstable nature of their endogenous counterparts and are likewise predominantly degraded by the proteasome. We concluded that the established reporter cell lines were suitable for subsequent genetic screening to identify IRF1 and cMYC stability regulators and that the established cell lines could be used for parallel screens to determine specificity of identified factors.

### Identification of the Cullin substrate receptor SPOP as a regulator of IRF1 abundance

The characterized mCherry-IRF1 and mCherry-cMYC reporter cell lines were subsequently used for genome-wide genetic screening for regulators of their protein abundance. To this end, a genome-wide lentiviral sgRNA library was integrated in the cell lines at a low multiplicity of infection (MOI) ensuring the targeting of one gene per cell (Fig. 2A, Appendix Fig. 1), followed by selection for G418-resistance to enrich for sgRNA-expressing cells (Appendix Fig. 2). At three and six days after Cas9 induction with Dox, the top 1-3% of cells with the highest and lowest mCherry levels were collected by flow cytometry (Fig. 2A, Appendix Fig. 3 and 4), to enrich for cells with decreased or increased levels of the unstable mCherry-IRF1 or mCherry-cMYC. In parallel, the top 1-3% of cells with the highest and lowest levels of the stable BFP control were collected to identify non-specific effects (Fig. 2A, Appendix Fig. 3 and 4).

**Figure 2.**
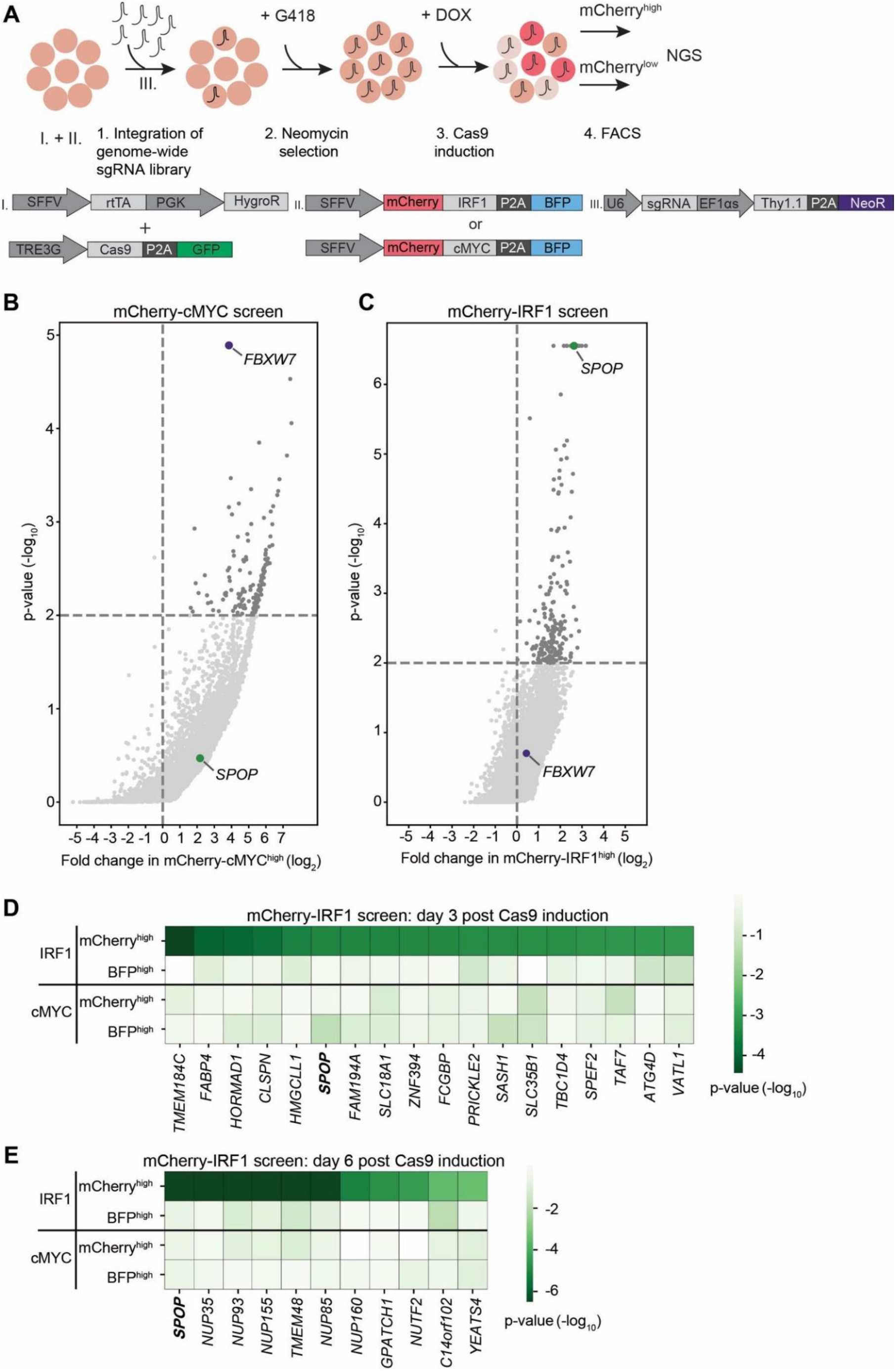
Identification of SPOP as a regulator of IRF1 abundance. **A)** Overview of FACS-based CRISPR-Cas9 knockout screening procedure using the RKO-mCherry-IRF1-P2A-BFP or RKO-mCherry-cMYC-P2A-BFP cell lines. Cells expressing high and low levels of mCherry-cMYC or mCherry-IRF1 protein were sorted and their integrated sgRNA coding sequences determined by Next Generation Sequencing. **B-C)** Read counts per million in the mCherry-cMYC^high^ and mCherry-IRF1^high^ cells at 6 days after Cas9 induction were compared to those in unsorted cells from the same day, sgRNA enrichment calculated by MAGeCK analysis, and log_2_-fold change and adjusted p-value plotted. Specific top hits SPOP and FBXW7 are shown for the **B)** mCherry-cMYC, or **C)** mCherry-IRF1 screen. **D-E**) 20 genes enriched in the mCherry-IRF1^high^ sorted populations with highest adjusted p-values were selected and any genes enriched in BFP^high^ or mCherry^low^ populations were excluded. Screen-specific adj. p-value heatmaps were generated for mCherry-IRF1^high^ cells at **D)** 3 days, or **E)** 6 days post-Cas9 induction.

The two harvesting time points were previously empirically determined to be optimal for capturing essential genes (day 3), and genes that encode proteins whose levels require longer times to deplete (day 6) (Fig. S1E-F) (de Almeida et al., 2021). Integrated sgRNA coding sequences from the selected cell pools were quantified by next generation sequencing and compared to relative frequencies determined in unsorted cells representing the initial library (Appendix Fig. 5). The robustness and uniformity of the screens in both cell lines is underpinned by the fact that all sgRNA coding sequences in unsorted samples were covered by 100-1000 reads (Appendix Fig. 6).

As anticipated, the mCherry-cMYC screen strongly identified its well-known degrader FBXW7 in mCherry-cMYC^high^ cells (Fig. 2B, Fig. S2A-D). Moreover, it identified other UPS components reported to mediate cMYC degradation (Fig. S2A-D), such as proteasome components (PSMC5, PSMD4, PSMD11), the proteasome regulator AKIRIN2 (de Almeida et al., 2021), and the UPS-related ATPase VCP/p97 (Heidelberger et al., 2018). These factors were not enriched in the BFP^high^ stable control cells (Fig. S2C-D), attesting to the specificity and identification power of the experimental system.

The mCherry-IRF1 screen identified 27 high confidence factors which specifically affected mCherry-IRF1 abundance, but neither mCherry-cMYC, nor the stable BFP controls (Fig. 2C-E). The strongest candidate enriched at both three and six days after Cas9 induction was the Cullin E3 ligase receptor Speckle Type BTB/POZ Protein (SPOP), which did not score in the mCherry-cMYC screen (Fig. 2B-E, Fig. S2E-F). In addition, seven nuclear transport-related factors (NUP35, NUP85, NUP93, NUP155, NUP160, TMEM48, NUTF2) were identified (Fig. 2E, Fig. S2F), suggesting that nuclear translocation of IRF1 or one of its degraders may be important for its degradation. Consistent with this notion, SPOP has been previously reported to localize to the nucleus and degrade nuclear substrates (Marzahn et al., 2016; Usher et al., 2021). Moreover, a chromatin reader component of the NuA4 histone acetyltransferase (HAT) complex (YEATS4) and two factors with limited functional annotation (GPATCH1, C14ORF102) were identified (Fig. 2E).

Previously described regulators of IRF1 proteasomal turnover such as FBXW7, MDM2, or STUB1/CHIP (Garvin et al., 2019, p. 1; Landré et al., 2013; Narayan, Pion, et al., 2011) did not score as significant IRF1 regulators in our screen, indicating that they are unlikely to play a significant role in IRF1 degradation in this cellular context. Importantly, FBXW7 was identified as the strongest factor mediating cMYC degradation (Fig. 2B, Fig. S2A-D), underpinning the functionality and specificity of the screening approach. In summary, our genetic screen identified the E3 ligase substrate receptor SPOP as a specific candidate to regulate IRF1 protein abundance.

### SPOP regulates IRF1 protein levels

To determine whether our genetic screen results were predictive, individual genes identified in cell pools with increased mCherry-cMYC or mCherry-IRF1 levels were targeted. Ablation of top-scoring putative regulators of cMYC (*FBXW7*, *UBE3C*, *VCP*) substantially increased mCherry-cMYC levels (Fig. S3A-C), whereas targeting SPOP specifically increased the abundance of mCherry-IRF1 (Fig. 3A, Fig. S3D).

**Figure 3.**
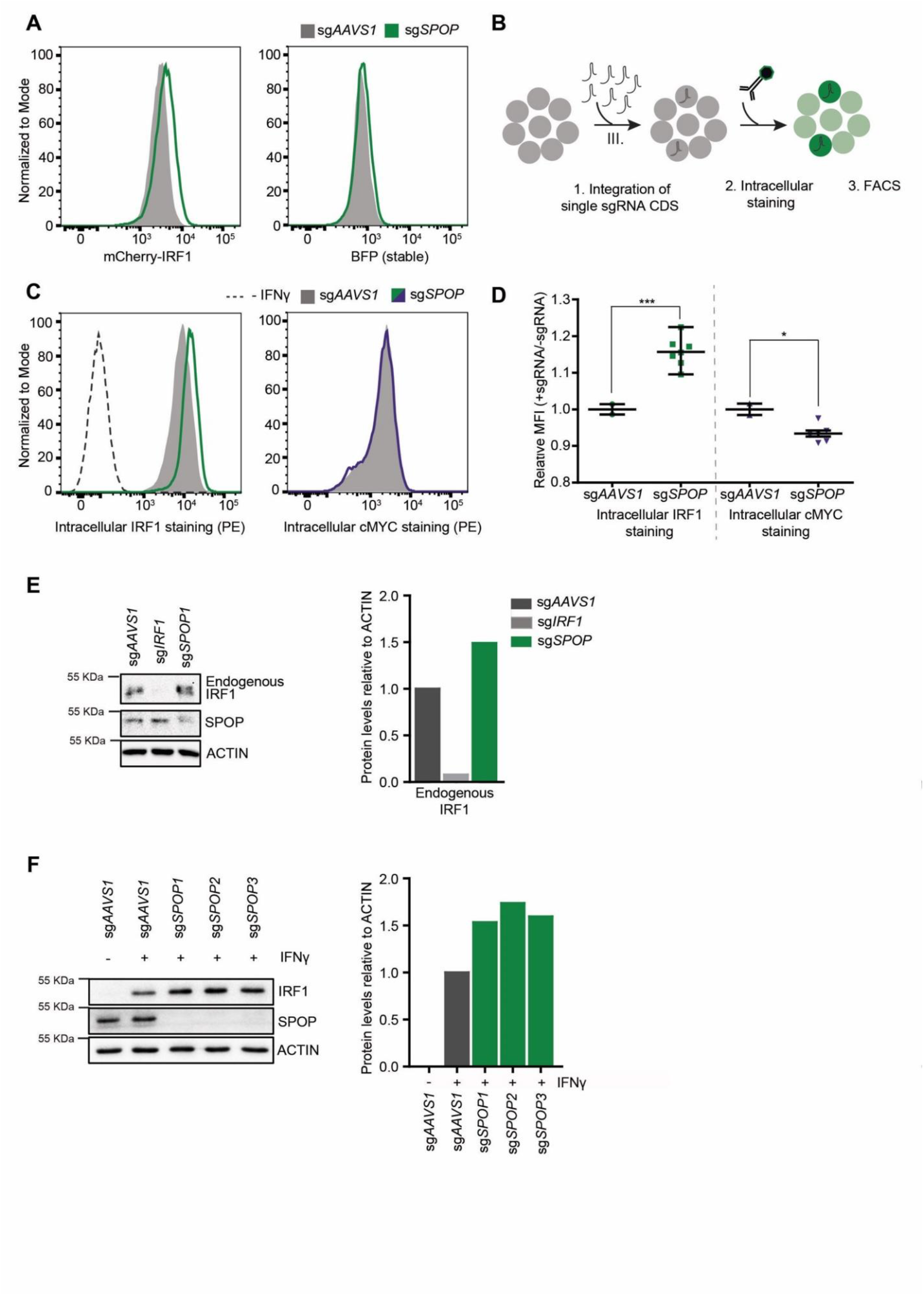
SPOP regulates IRF1 protein levels. **A)** RKO-mCherry-IRF1-P2A-BFP cells expressing sg*AAVS1* or sg*SPOP* were treated with Dox to induce Cas9 and subsequently mCherry-IRF1 and BFP protein levels were analyzed by flow cytometry. **B)** RKO-Cas9-P2A-GFP cells expressing sg*AAVS1* or sg*SPOP* were treated with Dox to induce Cas9, treated with IFNγ for 4 h., after which endogenous IRF1 or cMYC levels were determined by **C-D)** intracellular staining and flow cytometry, or **E)** Western blot. In **D)**, each data point represents targeting with a different, independent sg*SPOP* sequence; n = 2, unpaired two-tailed t-test. * p < 0.05, *** p < 0.001. **F)** Independent monoclonal RKO-Cas9-GFP cells lines with wt *IRF1* alleles derived from sg*AAVS1* expressing cells, or homozygous *SPOP* disrupted alleles were stimulated with IFNγ for 4 h., after which their IRF1 and SPOP protein levels were determined by Western blot, and **G)** quantified.

Next, we set out to assess whether SPOP regulates endogenous IRF1 protein levels. To this end, cells were transduced with sg*SPOP* expression vectors (Fig. 3B), Cas9 was induced with Dox, and their endogenous intracellular IRF1 and cMYC protein levels determined by flow cytometry (Fig. 3B-D). Consistent with our screen data (Fig. 2, Fig. S2), *SPOP* ablation increased endogenous IRF1 levels by 40%, whereas endogenous cMYC protein levels were not affected (Fig. 3C). In line with these results, *SPOP* ablation increased endogenous IRF1 abundance measured by Western blot (Fig. 3E).

The sgRNA library design with coverage of each gene by six sgRNAs made it unlikely that off-target effects of the *SPOP*-targeting sgRNAs were responsible for the selected phenotype of increased mCherry-IRF1. However, to measure whether individual sgRNAs would result in the same IRF1-stabilizing phenotype, we transduced cells individually with seven different, independent, sg*SPOP* sequences, and analyzed intracellular IRF1 levels by flow cytometry (Fig. 3D). All seven different sgRNAs significantly increased IRF1 levels compared to the two independent controls, indicating that specific *SPOP*-targeting underlies this phenotype, and not off-target effects (Fig. 3D). Consistent with this notion, Western blot analysis of three independent monoclonal *SPOP* knock-out cell lines showed that IRF1 levels were increased by 50-70% (Fig. 3F). This effect of *SPOP* ablation specifically affected IRF1 protein levels, as *IRF1* mRNA concentrations were not significantly changed in *SPOP* knock-out cells (Fig. S3E-F). In summary, these findings demonstrate that SPOP is important for regulating IRF1 protein levels, without affecting its mRNA concentrations.

### SPOP targets S/T-rich degrons in IRF1

Having identified SPOP in a phenotypic genetic screen, it was possible that the effects of *SPOP* ablation on IRF1 were direct, or indirect through unknown other mediators. Previous studies had identified an S/T-rich SPOP binding consensus of [Φ]-[Π]-[S]-[S/T]-[S/T], in which Φ and Π are non-polar and polar residues, respectively (Zhuang et al., 2009). We reasoned that the presence of any sequences matching this SPOP degron in IRF1 could mediate SPOP binding to IRF1, thereby targeting it for ubiquitination and degradation.

A multiple sequence alignment of IRF1 across species (Fig. S4A), identified four sequences in IRF1 matching the SPOP degron consensus (Fig. 4A), consistent with predictions (ScanProsite – Expasy, and (Hou et al., 2022). One of the putative SPOP degrons was identified in the region between the DNA-binding domain and the IAD2 domain, whereas the remaining three were predicted to be inside the IAD2 domain, which is important for transactivation and dimerization (Fig. 4A, Fig. S4A).

**Figure 4.**
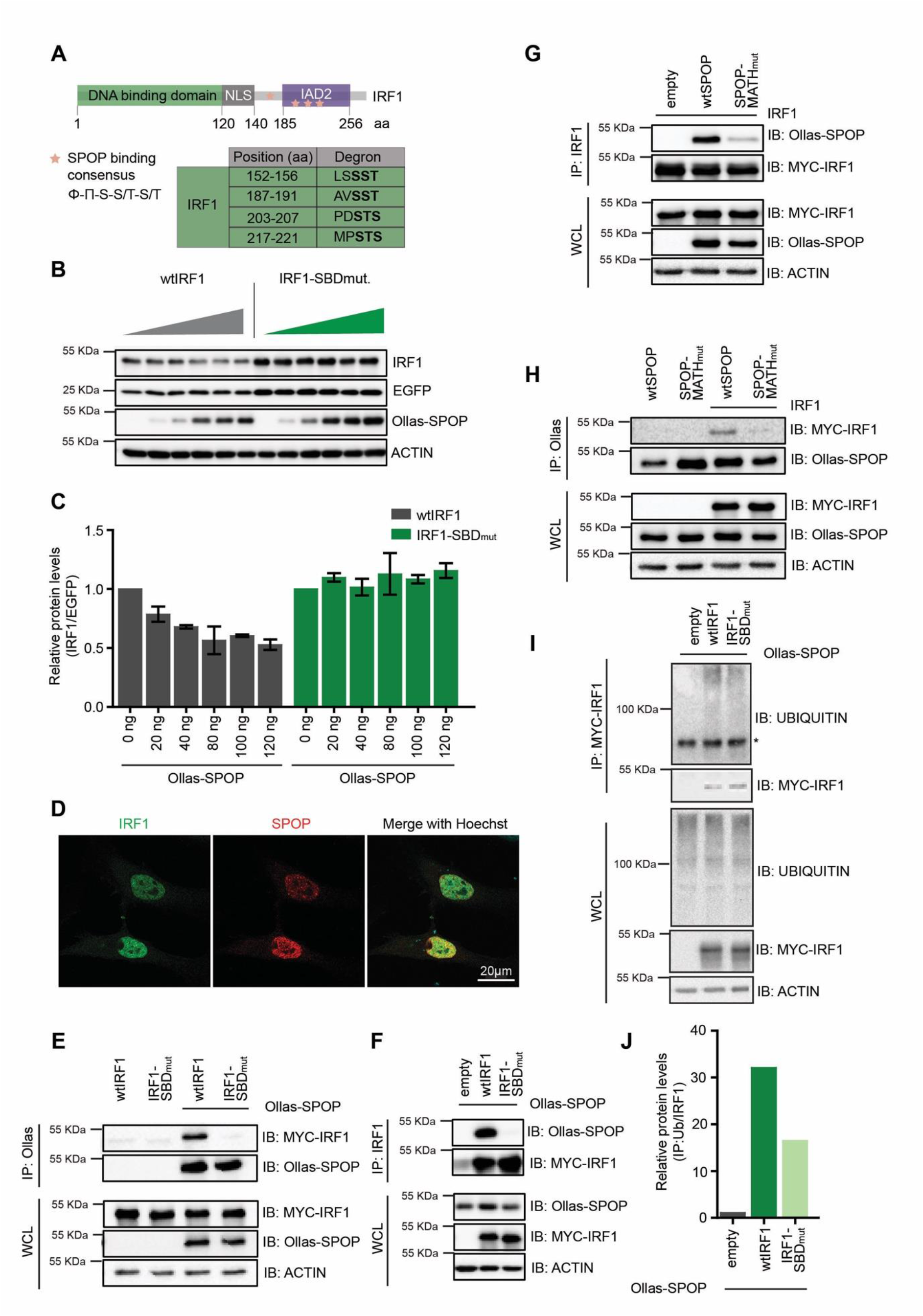
SPOP targets S/T-rich degrons in IRF1. **A)** Schematic representation of human IRF1 and its conserved domains. The SPOP consensus binding sequence is displayed and matching degron sequences in human IRF1 indicated. **B)** HEK-293T cells were co-transfected with expression plasmids for EGFP, Ollas-tagged wtSPOP, and either wtIRF1, or an IRF1 mutant in which the S/T residues in the four indicated SPOP Binding Domains (SBD) were mutated to A (IRF1-SBD_mut_). At 36 h. post-transfection, cell lysates were analyzed by Western blot and **C)** quantified. **D)** HeLa cells were transfected with expression plasmids for MYC-tagged IRF1 and FLAG-tagged SPOP. At 36 h. post-transfection, cells were fixed and localization analyzed by confocal immunofluorescence microscopy. **E-H)** HEK-293T cells were transfected with the indicated expression plasmids for wtIRF1, IRF1-SBD_mut_, Ollas-tagged wtSPOP, or a SPOP mutant in which its substrate interaction site in its MATH domain was mutated (SPOP-MATH_mut_). At 36 h. post-transfection, interaction was analyzed by co-IP. **I-J)** HEK-293T cells were transfected with the indicated expression plasmids for MYC-tagged wtIRF1 or IRF1-SBD_mut_. Subsequently, MYC-tagged IRF1 was immunoprecipitated and **I)** ubiquitination analyzed by Western blot and **J)** quantified. * indicates a non-specific product.

To test whether these predicted degron sites in IRF1 are important for SPOP-dependent degradation, an IRF1 mutant was generated in which the core S/T residues in each putative SPOP Binding Domain (SBD) were mutated to A residues (IRF1-SBD_mut_). Subsequently, wild-type IRF1 or IRF1-SBD_mut_ were expressed in the presence of increasing amounts of a wild-type SPOP expression plasmid (Fig. 4B). SPOP expression decreased wtIRF1 protein levels in a dose-dependent manner (Fig. 4B-C), without affecting the stable EGFP internal control. In contrast, protein levels of the IRF1 degron mutant remained unaffected by exogenous SPOP expression, suggesting that the four predicted degrons collectively are required for SPOP-dependent IRF1 degradation.

Next, we set out to investigate whether IRF1 and SPOP interact in cells and whether this interaction is mediated through the predicted IRF1 S/T-degrons. First, we analyzed whether IRF1 and SPOP reside in the same sub-cellular compartment. To this end, cells expressing exogenous IRF1 and SPOP were analyzed by immunofluorescence microscopy. Consistent with previous reports, both IRF1 and SPOP localized in the nucleoplasm (Fig. 4D), indicating that they are present in the same sub-cellular compartment, and thus could interact. However, given their disperse nucleoplasmic localization, co-localization could not be effectively addressed.

To directly assess complex formation between SPOP and IRF1, SPOP was immunoprecipitated from cells expressing wtIRF1 or IRF1-SBD_mut_. Subsequent Western blot analysis for IRF1 identified wtIRF1, but not IRF1-SBD_mut_ (Fig. 4E). The reciprocal approach in which wtIRF1 or IRF1-SBD_mut_ were immunoprecipitated, exclusively identified SPOP upon purification of wtIRF1, but not its degron mutant (Fig. 4F).

SPOP interaction with S/T degrons has been previously mapped to its MATH domain (Zhuang et al., 2009). In line with this observation, wtIRF1 interacted with wtSPOP, yet failed to co-immunoprecipitate MATH-domain mutated SPOP (SPOP-MATH_mut_) (Fig. 4G). Consistently, the reciprocal pull-down of wtSPOP identified wtIRF1, whereas the SPOP mutant showed strongly reduced interaction with IRF1 (Fig. 4H). In line with the four identified degrons being important for SPOP-dependent IRF1 turnover, IRF1 ubiquitination was reduced by 50% upon mutation of the SPOP binding sequences (Fig. 4I-J). From this, we concluded that the SPOP degrons are required for IRF1 poly-ubiquitination, although the remaining ubiquitination in the IRF1 mutant indicates that additional E3 ligases may ubiquitinate IRF1 as well. It remains to be determined whether these SPOP-independent poly-ubiquitin chains have a topology associated with proteasomal degradation.

Lastly, we tested whether wtSPOP or SPOP-MATH_mut_ could rescue IRF1 degradation in *SPOP* KO cells. To this end, expression plasmids for wtSPOP or its MATH domain mutant counterpart were delivered to *SPOP* KO cells, and endogenous IRF1 levels measured by flow cytometry (Fig. S4B). In line with previous observations, knockout of *SPOP* increased IRF1 protein levels (Fig. S4C). Expression of wtSPOP in *AAVS1*-targeted RKO cells did not alter endogenous IRF1 protein concentrations compared to an empty vector control (Fig. S4D, left panel), indicating that SPOP levels in these cells were likely not limiting for IRF1 turnover. In contrast, the elevated IRF1 protein levels in *SPOP* KO cells were decreased by exogenous wtSPOP expression to levels measured in wild-type cells, whereas they were not affected by the empty vector control (Fig. S4D, middle panel). Consistent with our interaction data and requirement of the SPOP MATH domain for IRF1 targeting (Fig. 4E-H), IRF1 levels were only rescued by wtSPOP expression, but not by its MATH domain mutant counterpart (Fig. S4D, right panel, and S4E).

Taken together, these data indicate that IRF1 is targeted by SPOP for ubiquitination through interaction of the SPOP MATH domain with S/T-rich degrons in IRF1, resulting in subsequent IRF1 degradation.

### SPOP controls IRF1-dependent transcriptional output

IRF1 is required for efficient transcription of interferon induced genes (ISGs). IRF1 is in particular important for transcriptional induction of a subset of ISGs, such as members of the guanylate binding protein (GBP) family in response to interferons (Ramsauer et al., 2007). By querying the harmonizome project for IRF1 transcriptional targets (Rouillard et al., 2016), we identified the ISGs *GBP2*, *GBP3*, *IFIT3*, and *NLRC5* as putatively IRF1-dependent immune effector ISGs, and *PARP14*, *APOL2*, and *DTX3L* as IRF1-dependent non-immune effector genes in RKO cells. *IRF1* itself and *STAT1* were included as IRF1-independent genes. We asked whether SPOP-dependent control of IRF1 protein abundance influences IRF1-dependent transcriptional output. To this end, mRNA levels of IRF1-dependent and - independent targets were measured by RT-qPCR in *SPOP*-deficient cells.

*SPOP* ablation did not significantly change mRNA concentrations of *IRF1* or *STAT1*, the transcription of which is IRF1-independent (Fig. 5A). In contrast, *SPOP* ablation significantly increased the mRNA levels of IRF1-dependent immune-related targets *GBP2*, *GBP3*, *IFIT3*, and *NLRC5* (Fig. 5A). The effect of *SPOP* loss was particularly pronounced on the well-known IRF1-specific target GBP2, which increased by 40-fold in *SPOP* KO cells (Fig. 5A). Moreover, IRF1-dependent non-immune transcripts *APOL2*, *PARP14*, and *DTX3L* (Fig. 5B) were likewise increased in the absence of SPOP. Together, these results indicate that loss of SPOP, and the resulting elevated IRF1 protein concentrations, increases the transcription of IRF1-dependent response genes.

**Figure 5.**
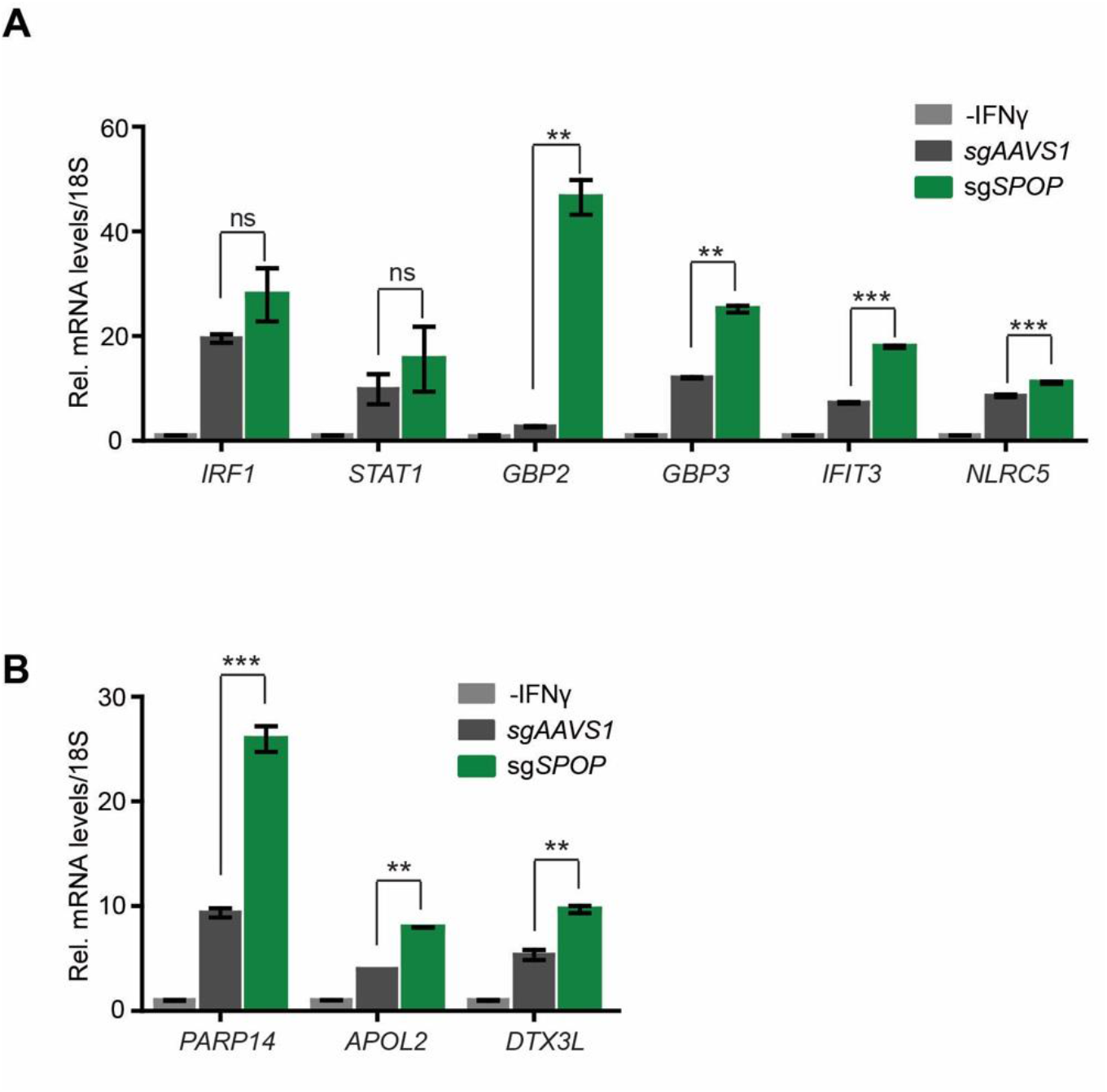
SPOP controls IRF1-dependent transcriptional output. Monoclonal RKO-mCherry-IRF1-P2A-BFP cells with either sg*AAVS1* or sg*SPOP* were stimulated with IFNγ for 4 h. and concentrations of the indicated **A)** immune-related, or **B)** non-immune-related mRNAs analyzed by RT-qPCR (n = 2).

## Discussion

The importance of IRF1 in myeloid cells for driving CXCL9 and CXCL10 expression and thereby being broadly critical for their innate immune response has been described (Forero et al., 2019). Moreover, IRF1 is a critical tumor suppressor, which is underpinned by the observation that IRF1 is mutated or lost in many cancers (Nozawa et al., 1999; Xie et al., 2003). Some ubiquitin E3 ligases controlling IRF1 protein turnover have been described, such as FBXW7, MDM2, and STUB1 (Garvin et al., 2019; Landré et al., 2013, p. 1; Narayan, Pion, et al., 2011). Their differential importance for regulating IRF1 turnover in different cell types and tissues draws an interesting functional parallel with cMYC, which is also targeted for degradation by cell-type specific E3 ligases (Sun et al., 2021).

Here we show that the Cullin E3 ligase receptor SPOP is important for IRF1 protein turnover. In contrast to previous findings in COS7 and MRC5 cells (Garvin et al., 2019), in our RKO cell-based experiments, FBXW7 did not contribute to IRF1 degradation, as its ablation did not alter IRF1 protein concentrations. Nevertheless, FBXW7 cellular functionality and efficient targeting was demonstrated, since it was key to cMYC degradation, consistent with previous reports (Welcker et al., 2004; Yada et al., 2004). These findings expand our knowledge on the pathways that can be employed by different cells to balance IRF1 levels. Moreover, it underpins the notion that, as for cMYC, the factors mediating IRF1 turnover can be cell-context-specific.

SPOP has been found to be differentially expressed in many cancers (Cuneo & Mittag, 2019; Guo et al., 2016; X.-M. Li et al., 2022; Patel et al., 2020; Song et al., 2020; Zhao et al., 2016). Moreover, mutations accumulate in SPOP target-recognition domains in endometrial and prostate cancers, altering their sensitivity to BET-inhibitor therapy (Dai et al., 2017; Janouskova et al., 2017; P. Zhang et al., 2017). Interestingly, these mutations enhanced BRD protein degradation in endometrial cancers, whereas it did the opposite in prostate cancers (Dai et al., 2017; Janouskova et al., 2017; P. Zhang et al., 2017). Thus, SPOP substrate targeting, its importance as a tumor-promoting or -repressing factor, and therapy sensitivity may be cancer- or cell-type specific. This exemplifies the cell context-dependent similarities between IRF1 and SPOP.

Several SPOP substrates important in a cancer context have been previously identified, such as BRD proteins, DRAK1 and LATS1 (Pang et al., 2022; Wang et al., 2020). Our data presented here adds IRF1 to this growing list of SPOP substrates. Since IRF1 levels are low in many cells in the absence of cytokine stimulation, for that reason it may have previously been missed as a SPOP interactor and substrate in unbiased proteomics efforts (Dai et al., 2017; Janouskova et al., 2017; P. Zhang et al., 2017). Thus, while the contributions of SPOP to cancer establishment or progression are likely through deregulation of a manifold of its substrates, our work suggests that IRF1 may be one of them. In line with our finding that SPOP could be critical for lowering the tumor-suppressive levels of IRF1, SPOP is overexpressed in most clear-cell renal cell carcinomas (ccRCC) (Patel et al., 2020; Zhao et al., 2016). The observation that IFNγ signaling and IRF1 expression are important biomarkers in ccRCC (Kong et al., 2022), addressing the importance of the SPOP-IRF1 axis in this cancer type may be of particular interest in future studies.

The complexity of IRF1 turnover regulation in different cell types is further illustrated by the fact that SPOP may influence IRF1 stability in myeloid cells in an opposite manner as we identified here in stromal cell types (Tawaratsumida et al., 2022). Recent work identified SPOP in myeloid cells as a key bridging molecule, indirectly repressing IRF1 phosphorylation, and its subsequent degradation. In line with this model, loss of *SPOP* in plasmacytoid dendritic cells and a mouse macrophage cell line decreased IRF1 (Tawaratsumida et al., 2022). This opposite SPOP phenotype and function from what we describe here could be explained by the fact that many non-immune cells do not express TLRs and their associated adapter proteins, which were shown to be required for SPOP complex formation and repression of IRF1 phosphorylation and degradation.

Importantly, IRF1 degradation in myeloid cells is seemingly dependent on active TLR-dependent immune signaling, whereas in the stromal, non-myeloid cell models used in this study, IRF1 degradation was independent of immune signaling. Several other studies have indicated that SPOP may inhibit TLR signaling through several different proposed means (Hu et al., 2021; Jin et al., 2020; Q. Li et al., 2020), suggesting that SPOP may balance biological output from the TLR-IRF1 axis through opposing effects. Together, these possible contrary effects of SPOP on IRF1 abundance in different cell types should thus be considered when addressing the impact of SPOP in the context of future whole-organism studies.

Previous work has indicated that SPOP recognizes nucleoplasmic targets with intrinsically disordered regions through multivalent interactions with S/T-rich degrons (Usher et al., 2021). In addition, its related paralog SPOP-like (SPOPL) has been proposed to regulate the oligomeric state of the SPOP E3 ligase complex (Errington et al., 2012). While our genetic screens indicate that SPOPL does likely not play an important role in IRF1 targeting, our data from this study are fully consistent with the reported substrate profile: IRF1 has IDRs and localizes almost exclusively to the nucleoplasm. Moreover, IRF1 is likely targeted by SPOP through recognition of multiple S/T-rich degrons, although we have not determined the contribution of the four individual S/T-rich degrons to IRF1 degradation.

The S/T residues in these SPOP binding domains are theoretical sites of phosphorylation, which as such could influence SPOP-IRF1 interaction. Indeed, previous work has indicated that SPOP preferentially binds some substrate S/T-rich degrons in their non-phosphorylated form, whereas in other substrates, phosphorylation of these sites seems a prerequisite for interaction and degradation (Jiang et al., 2021; Ostertag et al., 2019; Zhuang et al., 2009). Whether any of the identified IRF1 degrons are phosphorylated, and how this would affect its SPOP-dependent degradation, remains to be determined. IRF1 phosphorylation on S219 and S221 in the fourth degron was identified with low coverage in available IRF1 proteomics data (Garvin et al., 2019) and independently in a region that includes this site (Lin & Hiscott, 1999), suggesting that this particular identified SPOP binding domain could be post-translationally modified. The same sites were identified to be phosphorylated in breast cancer cells by the immune-activating kinase IKKε (Remoli et al., 2020). Phosphorylation under these conditions increased IRF1 proteasomal degradation. Since IKKε is activated under physiological conditions during infection, this could suggest that in some cellular contexts, ongoing immune signaling could thus provide a regulatory circuit driving transcription of *IRF1*, as well as providing its subsequent signal for degradation of the encoded IRF1 protein by SPOP.

Three of the identified S/T-rich degrons in IRF1 are located in its IAD2 transactivation domain. This domain is key for heterodimerization with other transcriptional activators or repressors (Antonczyk et al., 2019). Therefore, an additional regulatory implication of our findings could be that IRF1 heterodimerization through its IAD2 domain could shield three of the S/T-rich degrons from recognition by SPOP. As such, SPOP could thus preferentially target the inactive nucleoplasmic IRF1 pool that is not engaged in transcription.

Taken together, our study identified and characterized SPOP as a cellular factor targeting IRF1 for proteasomal degradation. It extends the growing set of E3 ligases which collectively mediate proper intracellular IRF1 protein levels in various different cellular contexts. Our findings lay the foundation for future studies to address how transcriptionally inactive and active immune-transcription factor pools are properly spatiotemporally targeted in diverse cell types.

## Materials and Methods

### Vectors

The lentiviral human genome-wide sgRNA library consists of 6 sgRNAs per gene for a whole coding human genome and was described elsewhere (Michlits et al., 2020). Lentiviral vectors expressing sgRNA from a U6 promoter and eBFP2 or iRFP from a PGK promoter have been described elsewhere (de Almeida et al., 2021). sgRNA CDSs were cloned in pLentiv2-U6-PGK-iRFP670-P2A-Neo (de Almeida et al., 2021) to perform knock-outs in RKO cell lines. The IRF1 and cMYC stability reporters (pLX-SFFV-mCherry-IRF1-P2A-eBFP2 and pLX-SFFV-mCherry-cMYC-P2A-eBFP2) were constructed by cloning the open reading frame (ORF) of human IRF1 or cMYC into a modified pLX303 vector (Addgene plasmid 25897). cDNAs encoding IRF1-SBD_mut_, wtSPOP and SPOP-MATH_mut_ were purchased from Twist Bioscience. The IRF1-SBD_mut_ degron mutant (S154A, S155A, T156A, S189A, S190A, T191A, S205A, T206A, S207A, S219A, T220A, S221A) and the SPOP-MATH_mut_ binding mutant (Y87C, F102C, W131G) were cloned into a modified pLX303 vector. The plasmids and sgRNAs used in this study are listed in the Appendix.

### Cell culture

RKO-mCherry-IRF1-P2A-eBFP2 or RKO-mCherry-cMYC-P2A-eBFP2 cells were generated by transducing RKO-Dox-Cas9 cells (de Almeida et al., 2021) with SFFV-MYC-mCherry-IRF1-P2A-eBFP2 or SFFV-MYC-mCherry-cMYC-P2A-eBFP2 lentiviral vectors. Cas9 was induced with 100 ng/ml of Doxycycline hyclate (Dox, Sigma-Aldrich, D9891) and mCherry-, eBFP2-, and GFP-positive single cells were sorted by FACS into 96-well plates using a FACSAria III cell sorter (BD Biosciences) to obtain single cell-derived clones. Cas9 genome editing and expression in the absence of Dox from the TRE3G promoter was tested in competitive proliferation assays. The cell lines and culture conditions used in this study are listed in the Appendix.

### Transfections

All transfections were performed by mixing DNA and polyethylenimine (PEI, Polysciences, 23966) in a 1:3 ratio (w/w) in DMEM (Sigma-Aldrich, D6429) without supplements. Transfection was performed using 500 ng of total DNA. The day before transfection, 1.5×10^5^ HEK-293T cells were seeded in 6-well clusters in fully supplemented DMEM media. Cells were harvested 36 h. after transfection, washed with ice cold PBS and stored at −80 °C until further processing.

### FACS-based CRISPR–Cas9 screens

Lentivirus-like particles were used to transduce RKO-mCherry-IRF1-P2A-eBFP2 or RKO-mCherry-cMYC-P2A-eBFP2 cells at a multiplicity of infection (MOI) of less than 0.2 TU/cell, and 500 to 1,000-fold library representation. The percentage of library-positive cells was determined after 3 days of transduction by immunostaining of the Thy1.1 surface marker and subsequent flow cytometric analysis. RKO cells with integrated lentiviral vectors were selected with G418 (1 mg/ml, Sigma-Aldrich, A1720) for 7 days. After G418 selection, 120 million cells of unsorted reference sample corresponding to 1,000-fold library representation were collected, and stored at −80 °C until further processing. Cas9 genome editing was induced with Dox (100 ng/ml, Sigma-Aldrich, D9891) and after 3 days and 6 days, cells were sorted by FACS. Cells were harvested, washed with PBS and stained with Fixable Viability Dye eFluor (1:1,000, eBioscience, 65-0865-14) for 30 min. Subsequently, cells were washed three times with PBS and sorted in supplemented RPMI-1640 using the FACSAria III cell sorter operated by BD FACSDiva software (v8.0). RKO cells were gated for non-debris, singlets, negative for Viability Dye, GFP-positive, mCherry-positive, and BFP-positive. 2-3% of cells with the lowest and 1-2% of cells with the highest mCherry or BFP signals were sorted into PBS. For both mCherry-IRF1 and mCherry-cMYC genome-wide screens, at least 1×10^6^ (mCherry^low^ and BFP^low^) and 2×10^6^ (mCherry^high^ and BFP^high^) cells were collected for each time point. Sorted samples were re-analyzed for purity, pelleted and stored at −80 °C until further processing. The gating strategy for flow cytometric cell sorting is shown in the Appendix.

### Protein half-life determination

To estimate endogenous IRF1 and cMYC as well mCherry-IRF1 and mCherry-cMYC protein half-lives, RKO mCherry-IRF1 or mCherry-cMYC expressing cells were treated with 30 μg/ml of cycloheximide (CHX, Sigma-Aldrich, C1988). At indicated time points, total protein extracts were generated using RIPA buffer, analyzed by Western blot, quantified, and normalized to ACTIN levels and to time point 0 as indicated. Single exponential decay curves were plotted using GraphPad Prism (v9), from which protein half-lives were calculated.

### Immunofluorescence confocal microscopy

200,000 HeLa cells/35-mm well were seeded onto coverslips (Marienfeld, 630-2190) and transfected using polyethylenimine transfection reagent (PEI, Polysciences, 23966) in a 1:6 (w/w) DNA/PEI ratio in non-supplemented DMEM with lentiviral plasmids expressing MYC-tagged wtIRF1 and FLAG-tagged wtSPOP. 48 h. after transfection, cells were fixed with 4% paraformaldehyde (PFA) for 15 min. Cells were permeabilized with 0.25% Triton X-100 in PBS for 5 min., followed by blocking of non-specific sites by incubation with 1% BSA for 30 min. at RT. Coverslips were incubated for 1 h. at RT with primary anti-IRF1 antibody, or anti-FLAG antibody in 1% BSA, followed by incubation with anti-rabbit IgG Alexa Fluor 488, or anti-mouse IgG DyLight 550 secondary antibodies, and incubation for 5 min. with 0.4X Hoechst in PBS. The coverslips were mounted using ProLong™ Gold Antifade Mountant (Invitrogen, P36934). Images were collected using a Zeiss LSM 980 confocal microscope at 40X magnification.

### Western blot

Cells were lysed in RIPA lysis buffer (50 mM Tris HCl (pH 7.4), 150 mM NaCl, 1% NP-40, 0.5% sodium Deoxycholate, 1 mM EDTA, 0.1% SDS, 0.1 mM PMSF, and 1X protease inhibitor cocktail). Cells were rotated for 20 min. at 4 °C and then centrifuged at 18,500 x g for 10 min. at 4 °C. Supernatants were transferred to new tubes and protein concentrations were determined by Pierce BCA Protein Assay Kit (Thermo Fisher Scientific, 23225). Between 20-50 μg of protein per sample was mixed with Laemmli sample buffer (62.5 mM Tris-HCl (pH 6.8), 5.8% glycerol, 2% SDS and 1.7% β-mercaptoethanol), and boiled for 5 min. Proteins were loaded on SDS polyacrylamide gels, the percentage of which was adjusted based on the molecular weight of the proteins of interest. Proteins were blotted on PVDF or nitrocellulose membranes at 4 °C for 1 h at 300 mA in Towbin buffer (25 mM Tris pH 8.3, 192 mM glycine and 20% ethanol). Membranes were blocked in 5% BSA in PBS-T for 1 h. at RT, and subsequently incubated with primary antibodies overnight at 4 °C. The next day, the membranes were washed three times for 5 min. each with PBS-T and incubated with HRP-coupled secondary antibodies for 1 h. at RT and imaged with the ChemiDoc™ Imaging System from Bio-Rad. Relative protein levels were quantified using Image Lab (Bio-Rad).

### Co-Immunoprecipitation assays

Cells were lysed in 100 μl of Frackelton lysis buffer (10 mM Tris (pH 7.4), 50 mM NaCl, 30 mM Na_4_P_2_O_7_, 50 mM NaF, 2 mM EDTA, 1% Triton X-100, 1 mM DTT, 0.1 mM PMSF, and 1X protease inhibitor cocktail). Cells were incubated on a rotating wheel at 4 °C for 30 min. and subsequently centrifuged at 20,000 x g at 4 °C for 30 min. The supernatant was transferred to a new tube and 10 μl (10% of the lysate used for the IPs) was collected as input. 300 μg of lysates were incubated overnight at 4 °C on a rotating wheel with an IgG Isotype Control (Cell Signaling Technology, 1:300), anti-Ollas (Novus, 1:100), or anti-IRF1 antibody (Cell Signaling Technology, 1:100). The next day, magnetic beads (Pierce Protein A/G Magnetic Beads, Thermo Fisher Scientific, 88803) used for anti-IRF1 antibody IPs, or Protein G Sepharose beads (Protein G Sepharose® 4 Fast Flow, Sigma-Aldrich, GE17-0618-01) used for anti-Ollas antibody IPs, were blocked by rotation in 3% BSA in Frackelton Buffer for 1 h. at 4 °C. 25 μl of beads were added to 300 μg of lysates and rotated for 2 h. at 4 °C. Then, the beads were washed five times with 1 ml of RIPA buffer, supplemented with 300 mM NaCl. Proteins were eluted by boiling in 2X disruption buffer (2.1 M urea, 667 mM β-mercaptoethanol and 1.4% SDS) for 5 min. at 95 °C.

### Immunoprecipitations for ubiquitination

Cells were lysed in 1 ml of RIPA lysis buffer (50 mM Tris-HCl (pH 7.4), 150 mM NaCl, 1% SDS, 0.5% sodium deoxycholate, 1% Triton X-100), supplemented with 40 mM N-Ethylmaleimide, 40 mM iodoacetamide, 25 U/ml Benzonase, and 1X protease inhibitor cocktail. Cells were incubated on a rotating wheel at 4 °C for 30 min., and centrifuged at 20,000 x g at 4 °C for 30 min. Supernatants were transferred to new tubes and 50 μl (20% of the lysates used for the IPs) were collected as input. 500 μg of lysates were incubated overnight at 4 °C on a rotating wheel with an IgG Isotype Control antibody (Cell Signaling Technology, 1:300), or anti-IRF1 antibody (Cell Signaling Technology, 1:100). The next day, magnetic beads (Pierce Protein A/G Magnetic Beads, Thermo Fisher Scientific, 88803) were blocked by rotation in 3% BSA in RIPA Buffer for 1 h. at 4 °C. 25 μl of beads were added to 500 μg of lysates and rotated for 2 h. at 4 °C. Subsequently, beads were washed five times with 1 ml of RIPA buffer, supplemented with 300 mM NaCl. Proteins were eluted by boiling in 2X disruption buffer (2.1 M urea, 667 mM β-mercaptoethanol and 1.4% SDS) for 5 min. at 95 °C.

### Lentivirus production and transduction

HEK-293T cells were transfected with DNA mixes containing lentiviral transfer plasmids, pCRV1-Gag-Pol (Hatziioannou et al., 2004) and pHCMV-VSV-G (Yee et al., 1994) using polyethylenimine (PEI, Polysciences, 23966) in a 1:3 μg DNA/PEI ratio in non-supplemented DMEM. Virus containing supernatants were clarified of cellular debris by filtration through a 0.45 μm filter. Virus-like particles were directly used after harvesting, or stored at −80 °C. Target cells were transduced in the presence of 5 μg/ml polybrene (Sigma-Aldrich, TR1003G).

### Intracellular staining for flow cytometry

For staining of intracellular proteins, cells were collected, washed twice with PBS and subsequently fixed with 2% PFA for 15 min. at RT. After one wash with FACS buffer (PBS supplemented with 1% FCS), cells were permeabilized in ice-cold MeOH for 10 min. and pelleted three times for 10 min. at 900 x g. Subsequently, cells were incubated for 10 min. at RT in FACS buffer to inhibit non-specific antibody binding. Cells were then incubated with primary antibodies, or left unstained for 1 h. at RT. Following two FACS buffer washes, cells were incubated with secondary antibodies for 15 min. at 4 °C., washed two times with FACS buffer and resuspended in FACS buffer for flow cytometric analysis on an LSRFortessa (BD Biosciences) operated by BD FACSDiva software (v8.0). FACS data were analyzed in FlowJo (v10.8).

### Next-generation sequencing library preparation

Next-generation sequencing (NGS) libraries of sorted and unsorted control samples were processed as previously described (de Almeida et al., 2021). In brief, isolated genomic DNA was subjected to two-step PCR. The first PCR allowed the amplification of the integrated sgRNA cassettes, the second PCR introduced the Illumina adapters. Purified PCR products’ size distribution and concentrations were measured using a fragment analyzer (Advanced Analytical Technologies). Equimolar ratios of the obtained libraries were pooled and sequenced on a HiSeq 2500 platform (Illumina). Primers used for library amplification are listed in the Appendix.

### Analysis of pooled CRISPR screens

The analysis of the CRISPR–Cas9 screens was carried out as previously described (de Almeida et al., 2021). In brief, sgRNAs enriched in day 3 and day 6 post-Cas9 induction-sorted samples were compared against the matching unsorted control populations harvested on the same days using MAGeCK (W. Li et al., 2014). In addition, depletion of sgRNAs over time was calculated by comparing the unsorted populations to the initial day 0 population.

### RNA isolation, cDNA synthesis, and qPCR

Total RNA was extracted from RKO-Dox-Cas9-P2A-GFP cells harboring *AAVS1-* or *SPOP*-targeting sgRNAs. 1×10^6^ cells were lysed using Trizol reagent (Thermo Fisher Scientific, 5596-018) and treated with Turbo DNase (Thermo Fisher Scientific, 10792877). cDNA was prepared using Oligo (dT18) Primer and RevertAid Reverse Transcriptase (Thermo Fisher Scientific, EP0441). Real-time PCR experiments were run on a Mastercycler (Biorad), using SYBR Green (Thermo Fisher Scientific). Primers for qPCR are listed in the Appendix.

## Acknowledgements

We thank Kitti Csalyi, Johanna Stranner, and Thomas Sauer at the Max Perutz Labs BioOptics FACS Facility for expert support, the Vienna Biocenter Core Facilities (VBCF) for Next Generation Sequencing analysis, and Aleksej Drino for help with plotting of NGS data. We are grateful to the ‘Signaling Mechanisms in Cellular Homeostasis’ doctoral program community, Manuela Baccarini, Pavel Kovarik and their labs for their technical expertise and help.

## Author contributions

M.V., I.S., A.C.G., M.M., and V.B. performed experiments and analyzed data. J.J., M.de A., and M.H. established critical reagents and methodology. J.W. and K.F. performed bioinformatic analyses. J.Z. and T.D. provided critical input on experimental designs and data analyses. G.A.V. designed and supervised the study, and wrote the manuscript, together with M.V. and I.S., with feedback from all co-authors.

---

**Figure S1.**
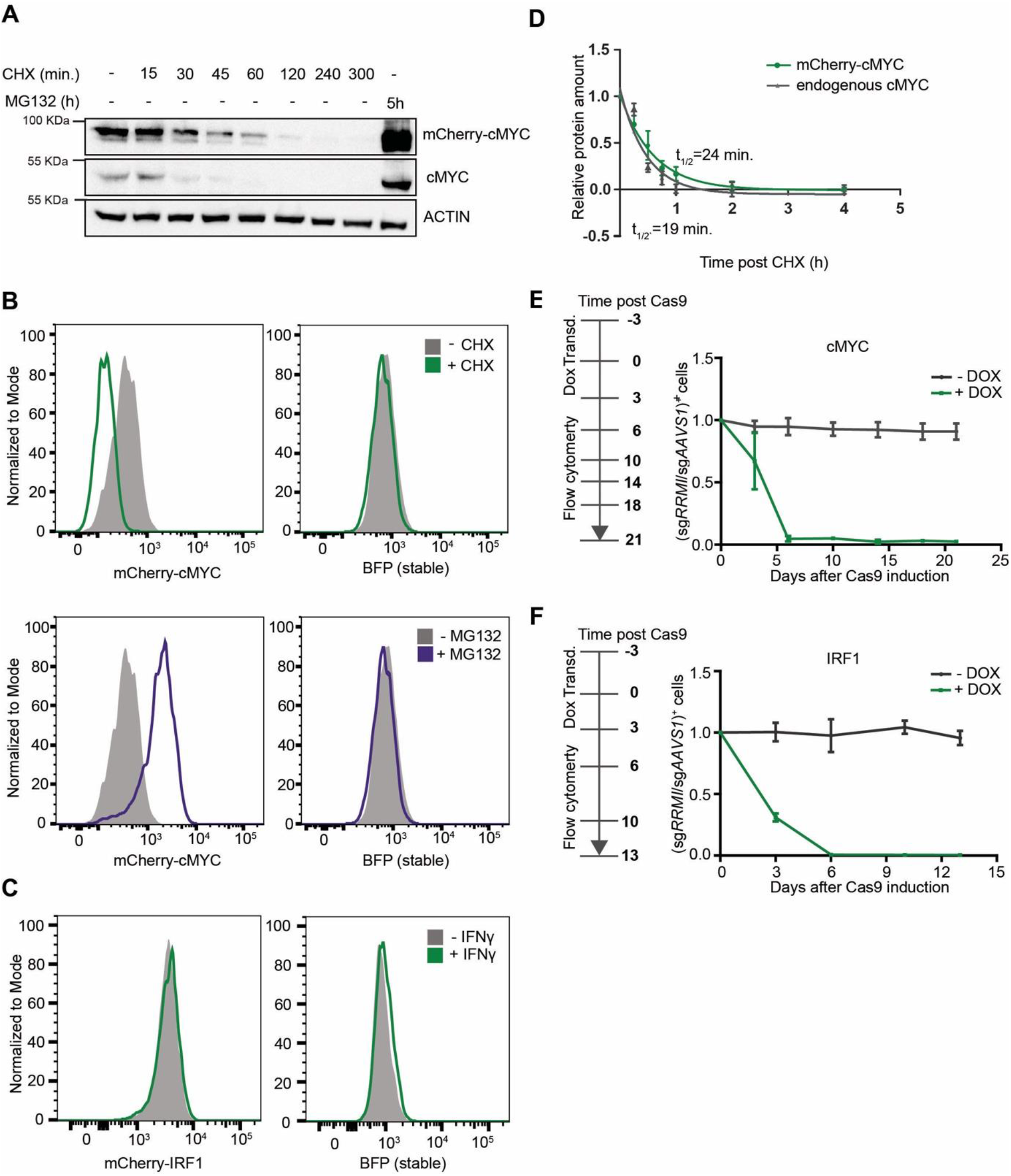
Establishment of an IRF1 protein abundance reporter cell line for genetic screening. **A)** RKO-mCherry-cMYC-P2A-BFP cells were incubated with CHX for the indicated times in the absence or presence of proteasome inhibitor MG132, after which mCherry-cMYC and endogenous cMYC levels were analyzed by Western blot. **B)** RKO-mCherry-cMYC-P2A-BFP cells were incubated with CHX or MG132 for 4 h., after which mCherry-cMYC and BFP protein levels were analyzed by flow cytometry. **C)** RKO-mCherry-IRF1-P2A-BFP cells were stimulated with IFNγ for 4 h., after which mCherry-IRF1 and BFP protein levels were determined by flow cytometry. **D)** RKO-mCherry-cMYC-P2A-BFP cells were incubated with CHX for the indicated times. Non-saturated Western blot signals for mCherry-cMYC and endogenous cMYC protein were quantified, normalized to ACTIN levels, plotted, and used for half-life calculation. n = 3. **E)** RKO-mCherry-cMYC-P2A-BFP cells, or **F)** RKO-mCherry-IRF1-P2A-BFP cells expressing Dox-inducible Cas9-P2A-GFP and sgRNAs targeting either *AAVS1*, or the essential gene *RRM1* were cultured in the presence or absence of Dox. At the indicated times, the relative abundances of sgRNA vector-positive cells were analyzed by flow cytometry and plotted. n = 2.

**Figure S2.**
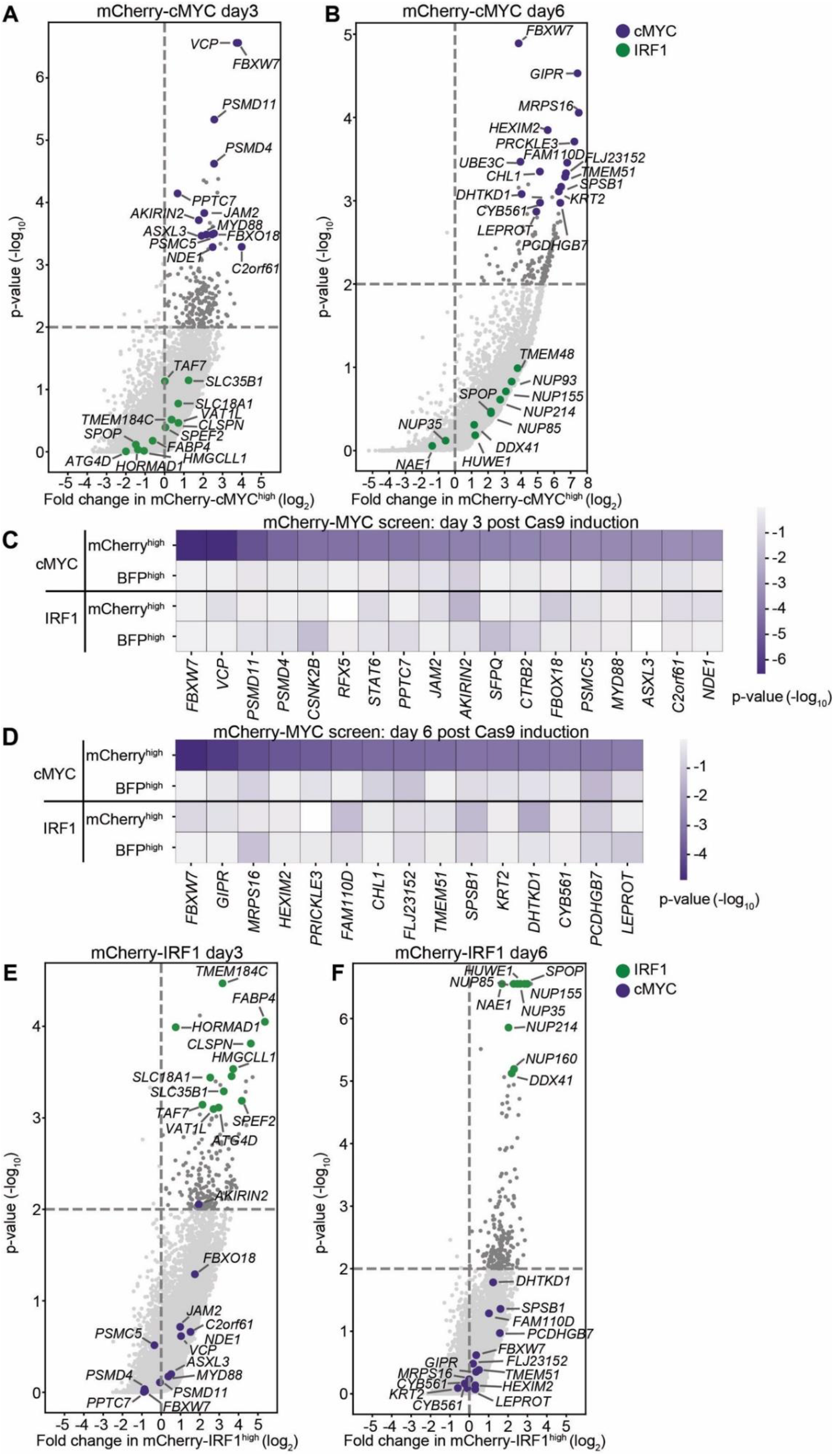
Identification of SPOP as a regulator of IRF1 abundance. **A-B)** Read counts per million in the mCherry-cMYC^high^ cells at **A)** 3 days post-Cas9 induction, and **B)** 6 days post-Cas9 induction were compared to those in unsorted cells from the same day, sgRNA enrichment calculated by MAGeCK analysis, and log_2_-fold change and adjusted p-value plotted. Specific top hits for the cMYC screen are show in purple, whereas IRF1 screen-specific genes are shown in green. **C-D**) 20 genes enriched in the mCherry-cMYC^high^ sorted populations with highest adjusted p-values were selected and any genes enriched in BFP^high^ or mCherry^low^ populations were excluded. Screen-specific adj. p-value heatmaps were generated for mCherry-cMYC^high^ cells at **C)** 3 days, or **D)** 6 days post-Cas9 induction. **E-F**) mCherry-IRF1 screen log_2_-fold change and adjusted p-values were plotted for mCherry-IRF1^high^ sorted cells at **E)** 3 days, or **F)** 6 days post-Cas9 induction. Specific top hits for the IRF1 screen are shown in green, whereas cMYC screen-specific genes are shown in purple.

**Figure S3.**
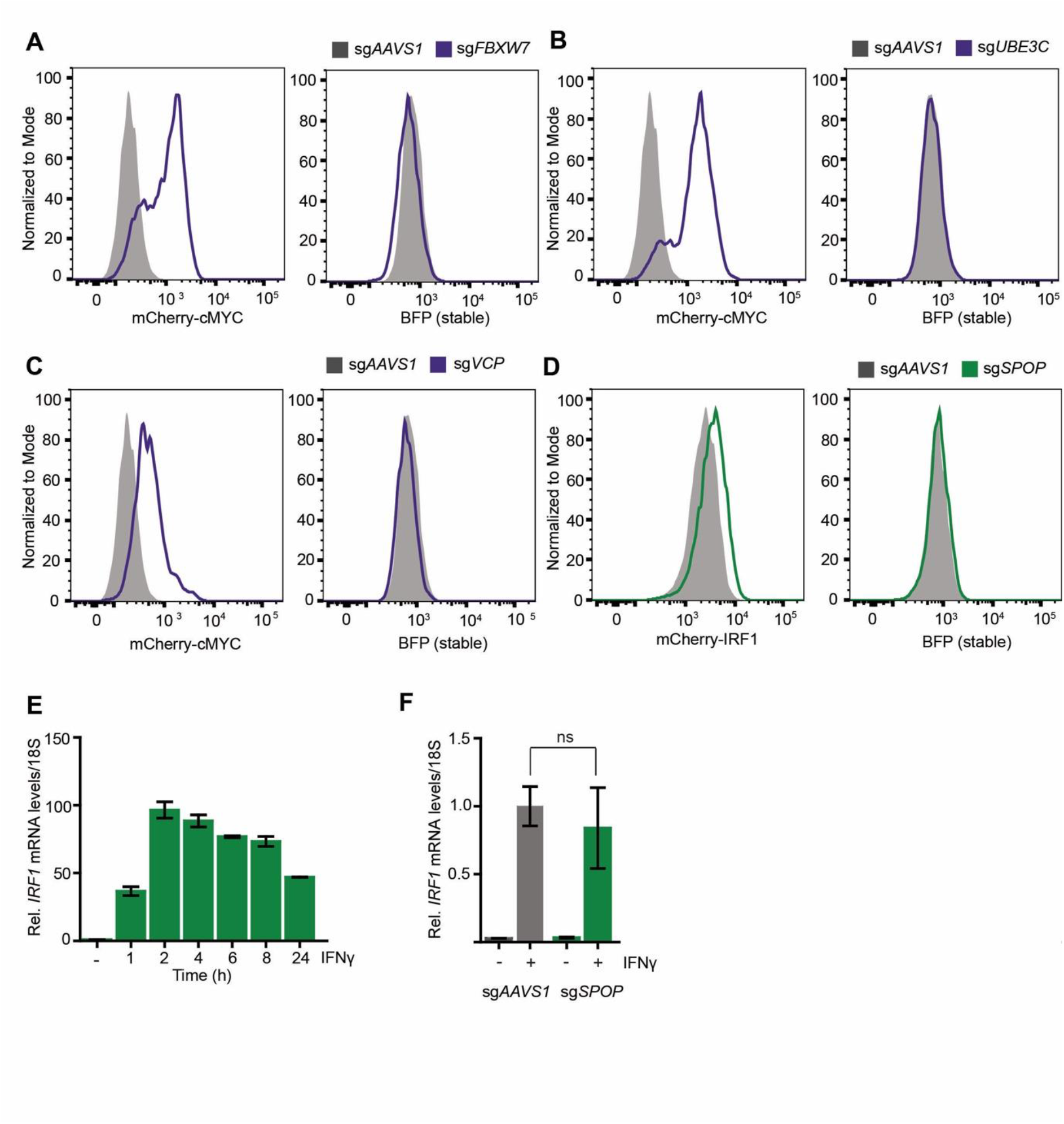
SPOP regulates IRF1 protein levels. **A-C)** RKO-mCherry-cMYC-P2A-BFP cells expressing sg*AAVS1* were compared with cells expressing **A)** sg*FBXW7*, **B)** sg*UBE3C*, or **C)** sg*VCP* after treatment with Dox for 6 (sg*FBXW7*, sg*UBE3C*), or 3 days (sg*VCP*) to induce Cas9 and subsequently mCherry-cMYC and BFP protein levels were analyzed by flow cytometry. **D)** RKO-mCherry-IRF1-P2A-BFP cells expressing sg*AAVS1* or sg*SPOP* were treated with Dox for 6 days to induce Cas9 and subsequently mCherry-IRF1 and BFP protein levels were analyzed by flow cytometry. **E)** RKO-Cas9-P2A-BFP cells were stimulated with IFNγ for the indicated times, after which *IRF1* mRNA levels were analyzed by RT-qPCR (n = 2). **F)** RKO-Cas9-P2A-BFP cells expressing either sg*AAVS* or sg*SPOP* were stimulated with IFNγ for 4 h. Subsequently, *IRF1* mRNA levels were analyzed by RT-qPCR (n = 2).

**Figure S4.**
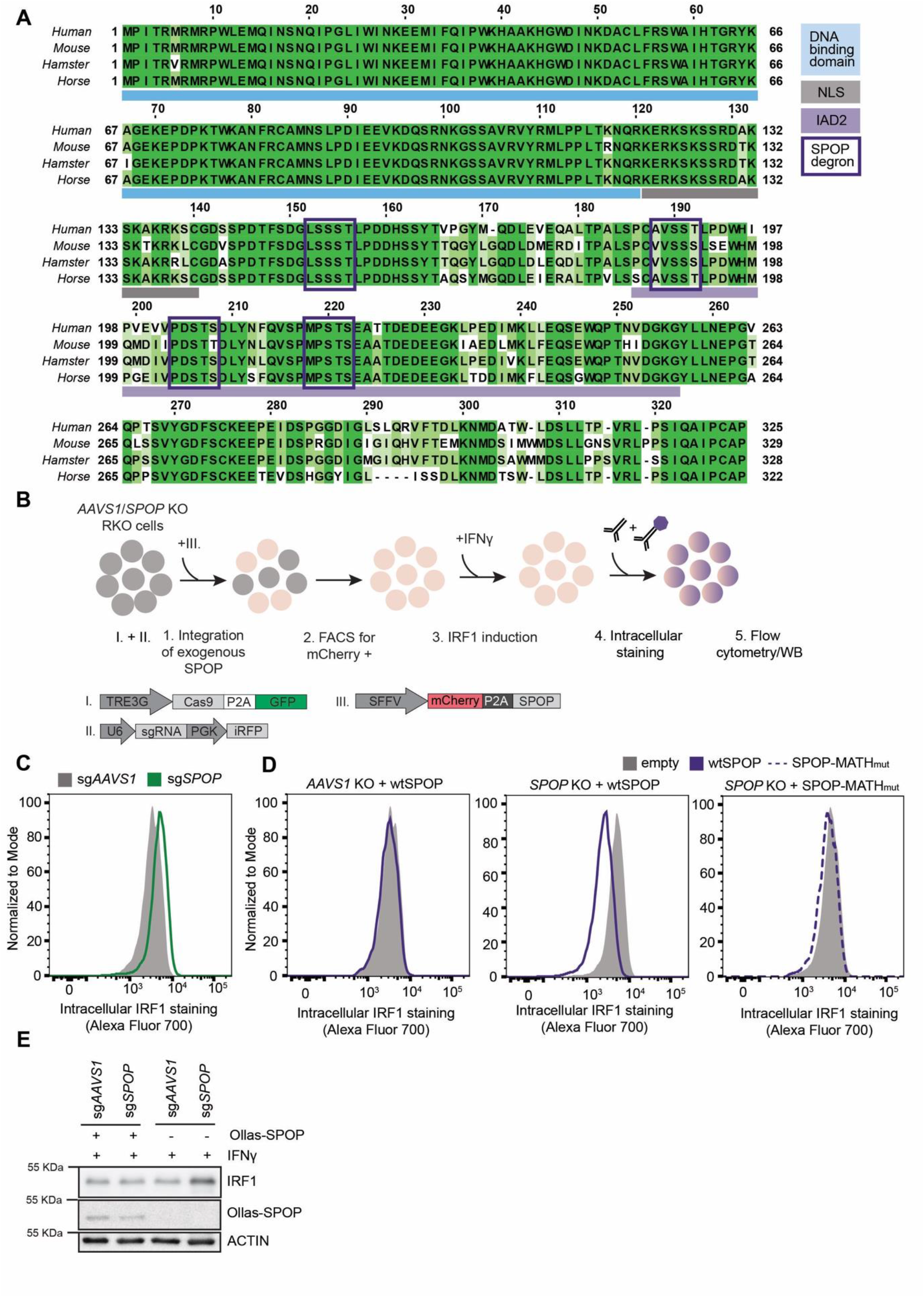
SPOP targets S/T-rich degrons in IRF1. **A)** IRF1 protein multiple sequence alignment. Predicted SPOP Binding Domains are indicated with boxes. **B)** Overview of rescue experiments. **C)** Endogenous IRF1 protein levels determined in wild-type (*AAVS1* KO), or *SPOP* KO RKO-Cas9-P2A-BFP cells by intracelluar staining and flow cytometry. **D-E)** Wild-type (*AAVS1* KO), or *SPOP* KO RKO-Cas9-P2A-BFP cells were transduced with vectors expressing either mCherry-P2A-wtSPOP, or mCherry-P2A-SPOP-MATH_mut_. Subsequently, endogenous IRF1 protein levels were determined by **D)** flow cytometry, or **E)** Western blot.

## Appendix

## Supplementary Materials and Methods

## Antibodies

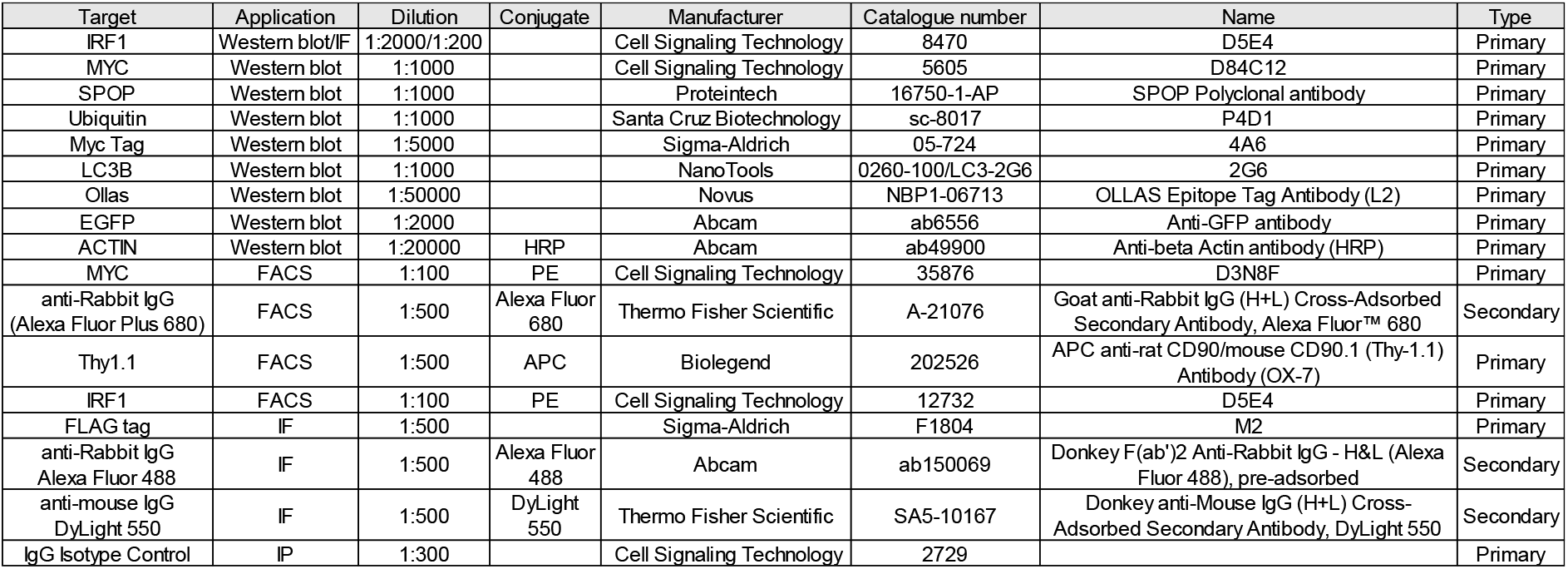

Supplementary Materials and Methods Antibodies:
| Target | Application | Dilution | Conjugate | Manufacturer | Catalogue number | Name | Type |
| --- | --- | --- | --- | --- | --- | --- | --- |
| IRF1 | Western blot/IF | 1:2000/1:200 |  | Cell Signaling Technology | 8470 | D5E4 | Primary |
| MYC | Western blot | 1:1000 |  | Cell Signaling Technology | 5605 | D84C12 | Primary |
| SPOP | Western blot | 1:1000 |  | Proteintech | 16750-1-AP | SPOP Polyclonal antibody | Primary |
| Ubiquitin | Western blot | 1:1000 |  | Santa Cruz Biotechnology | sc-8017 | P4D1 | Primary |
| Myc Tag | Western blot | 1:5000 |  | Sigma-Aldrich | 05-724 | 4A6 | Primary |
| LC3B | Western blot | 1:1000 |  | NanoTools | 0260-100/LC3-2G6 | 2G6 | Primary |
| Ollas | Western blot | 1:50000 |  | Novus | NBP1-06713 | OLLAS Epitope Tag Antibody (L2) | Primary |
| EGFP | Western blot | 1:2000 |  | Abcam | ab6556 | Anti-GFP antibody | Primary |
| ACTIN | Western blot | 1:20000 | HRP | Abcam | ab49900 | Anti-beta Actin antibody (HRP) | Primary |
| MYC | FACS | 1:100 | PE | Cell Signaling Technology | 35876 | D3N8F | Primary |
| anti-Rabbit IgG (Alexa Fluor Plus 680) | FACS | 1:500 | Alexa Fluor 680 | Thermo Fisher Scientific | A-21076 | Goat anti-Rabbit IgG (H+L) Cross-Adsorbed Secondary Antibody, Alexa Fluor™ 680 | Secondary |
| Thy1.1 | FACS | 1:500 | APC | Biolegend | 202526 | APC anti-rat CD90/mouse CD90.1 (Thy-1.1) Antibody (OX-7) | Primary |
| IRF1 | FACS | 1:100 | PE | Cell Signaling Technology | 12732 | D5E4 | Primary |
| FLAG tag | IF | 1:500 |  | Sigma-Aldrich | F1804 | M2 | Primary |
| anti-Rabbit IgG Alexa Fluor 488 | IF | 1:500 | Alexa Fluor 488 | Abcam | ab150069 | Donkey F(ab')<sub>2</sub> Anti-Rabbit IgG - H&L (Alexa Fluor 488), pre-adsorbed | Secondary |
| anti-mouse IgG DyLight 550 | IF | 1:500 | DyLight 550 | Thermo Fisher Scientific | SA5-10167 | Donkey anti-Mouse IgG (H+L) Cross-Adsorbed Secondary Antibody, DyLight 550 | Secondary |
| IgG isotype Control | IP | 1:300 |  | Cell Signaling Technology | 2729 |  | Primary |

## Reagents

Reagents:
| Description | Abbreviation | Application | Dilution/concentration | Manufacturer | Catalogue number |
| --- | --- | --- | --- | --- | --- |
| IFNγ human recombinant | IFNγ | Cell culture | 10 ng/mL | Enzo | ENZ-PRT141-0100 |
| Cycloheximide | CHX | Cell culture | 40 μg/mL | Sigma-Aldrich | C1988 |
| MG132 | MG132 | Cell culture | 10 μM | Sigma-Aldrich | M7449 |
| Epoxomicin | EPX | Cell culture | 10 μM | Gentaur Molecular Products | 607-A2606 |
| Bafilomycin A1 | Baf | Cell culture | 0.4 μM | InvivoGen | tlrl-baf1 |
| Doxycycline hyclate | DOX | Cell culture | 500 ng/mL | Sigma-Aldrich | D9891 |
| G418 disulfate salt | G418 | Cell culture | 0.5-1 mg/mL | Sigma-Aldrich | A1720 |
| Benzonase® Nuclease | | IP | 250 U/μl | Merck | 70746 |

## Plasmids

Plasmids:
| Plasmid | Purpose | Reference or source |
| --- | --- | --- |
| SFFV-pLX303-MYC-mCherry-IRF1-P2A-eBFP2 | IRF1 reporter | This study |
| SFFV-pLX303-MYC-mCherry-IRF1-SBDmut.-P2A-eBFP2 | IRF1 reporter | This study |
| SFFV-pLX303-MYC-mCherry-cMYC-P2A-eBFP2 | MYC reporter | This study |
| SFFV-pLX303-MYC-mCherry-P2A-Ollas-SPOP | SPOP reporter | This study |
| SFFV-pLX303-MYC-mCherry-P2A-Ollas-SPOP-MATHmut. | SPOP reporter | This study |
| pLentiv2-U6-PGK-iRFP670-P2A-Neo | single sgRNA expression | de Almeida M, Hinterndorfer M et al, Nature 2021 |
| pLentiv2-U6-PGK-eBFP2-P2A-Neo | single sgRNA expression | de Almeida M, Hinterndorfer M et al, Nature 2021 |
| PRRL-PBS-U6-sgRNA-EF1as-Thy1-P2A-NeoR (sgETN) | Library sgRNA | Johannes Zuber, IMP |
| SFFV-pLX303-MYC-IRF1 | Immunofluorescence | This study |
| SFFV-pLX303-FLAG-SPOP | Immunofluorescence | This study |

*In plasmid names, dashes denote separate elements, points indicate elements fused into a single open reading frame.*

## sgRNA sequences

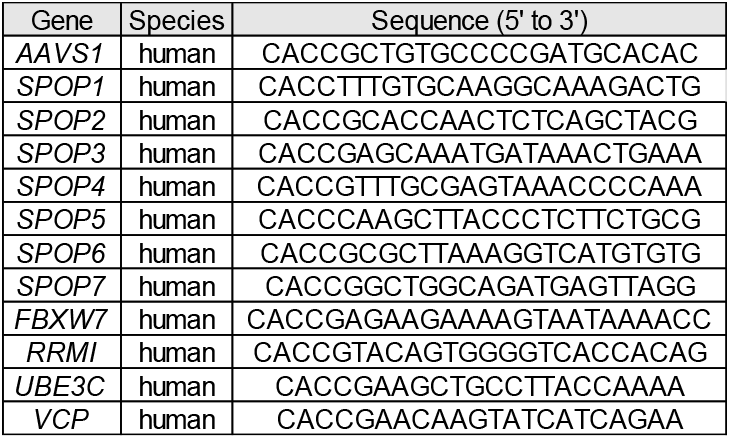

## Cell lines

Cell lines:
| Cell lines and primary cells | Type | Reference or source | Purpose | Media | Supplements |
| --- | --- | --- | --- | --- | --- |
| RKO | human rectal carcinoma | CRL-2577 | parental cell line | RPMI-1640 (Thermo Fisher Scientific, 21875) | 8.6% FBS (Sigma-Aldrich, F7524), L-glutamine (3.5 mM, Gibco), sodium pyruvate (1.7 mM, Sigma-Aldrich), and 0.9% penicillin/streptomycin (Sigma-Aldrich, P4333) |
| RKO | human colon carcinoma | de Almeida M, Hinterdorfer M et al, Nature 2021 | Dox-inducible Cas9 | RPMI 1640 (Thermo Fisher Scientific, 21875) | 8.6% FBS (Sigma-Aldrich, F7524), L-glutamine (3.5 mM, Gibco), sodium pyruvate (1.7 mM, Sigma-Aldrich), and 0.9% penicillin/streptomycin (Sigma-Aldrich, P4333) |
| RKO-dox-Cas9 | human rectal carcinoma | de Almeida M, Hinterdorfer M et al, Nature 2021 | Dox-inducible Cas9 | RPMI 1640 (Thermo Fisher Scientific, 21875) | 8.6% FBS (Sigma-Aldrich, F7524), L-glutamine (3.5 mM, Gibco), sodium pyruvate (1.7 mM, Sigma-Aldrich), and 0.9% penicillin/streptomycin (Sigma-Aldrich, P4333) |
| RKO-dox-Cas9 mCherry-IRF1-P2A-BFP | human rectal carcinoma | This study | mCherry-IRF1 reporter | RPMI 1640 (Thermo Fisher Scientific, 21875) | 8.6% FBS (Sigma-Aldrich, F7524), L-glutamine (3.5 mM, Gibco), sodium pyruvate (1.7 mM, Sigma-Aldrich), and 0.9% penicillin/streptomycin (Sigma-Aldrich, P4333) |
| RKO-dox-Cas9 mCherry-cMYC-P2A-BFP | human rectal carcinoma | This study | mCherry-cMYC reporter | RPMI 1640 (Thermo Fisher Scientific, 21875) | 8.6% FBS (Sigma-Aldrich, F7524), L-glutamine (3.5 mM, Gibco), sodium pyruvate (1.7 mM, Sigma-Aldrich), and 0.9% penicillin/streptomycin (Sigma-Aldrich, P4333) |
| HEK293T | human kidney neural tissue | CRL-3216 | IRF1-SBDmut. SPOP-MATHmut. Transfection | Dulbecco's modified Eagle's medium (DMEM; Sigma-Aldrich, D6429) | 8.6% FBS (Sigma-Aldrich, F7524) and 0.9% penicillin/streptomycin (Sigma-Aldrich, P4333) |
| Lenti-X 293T | human kidney neural tissue | Takara, Cat# 632180 | VLP production | Dulbecco's modified Eagle's medium (DMEM; Sigma-Aldrich, D6429) | 8.6% FBS (Sigma-Aldrich, F7524) and 0.9% penicillin/streptomycin (Sigma-Aldrich, P4333) |
| HeLa | Cervical adenocarcinoma | CCL-2 | Immunofluorescence microscopy | Dulbecco's modified Eagle's medium (DMEM; Sigma-Aldrich, D6429) | 8.6% FBS (Sigma-Aldrich, F7524) and 0.9% penicillin/streptomycin (Sigma-Aldrich, P4333) |

## qPCR primers

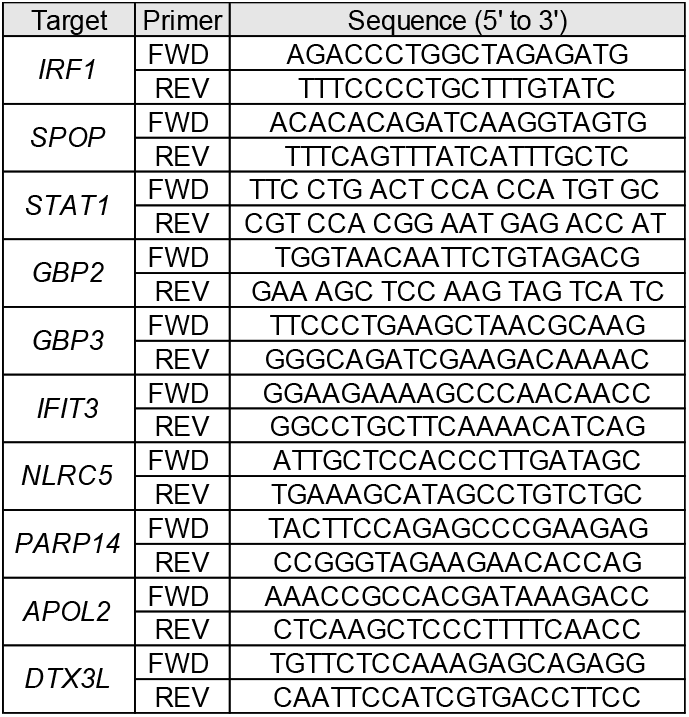

## NGS library PCR primers

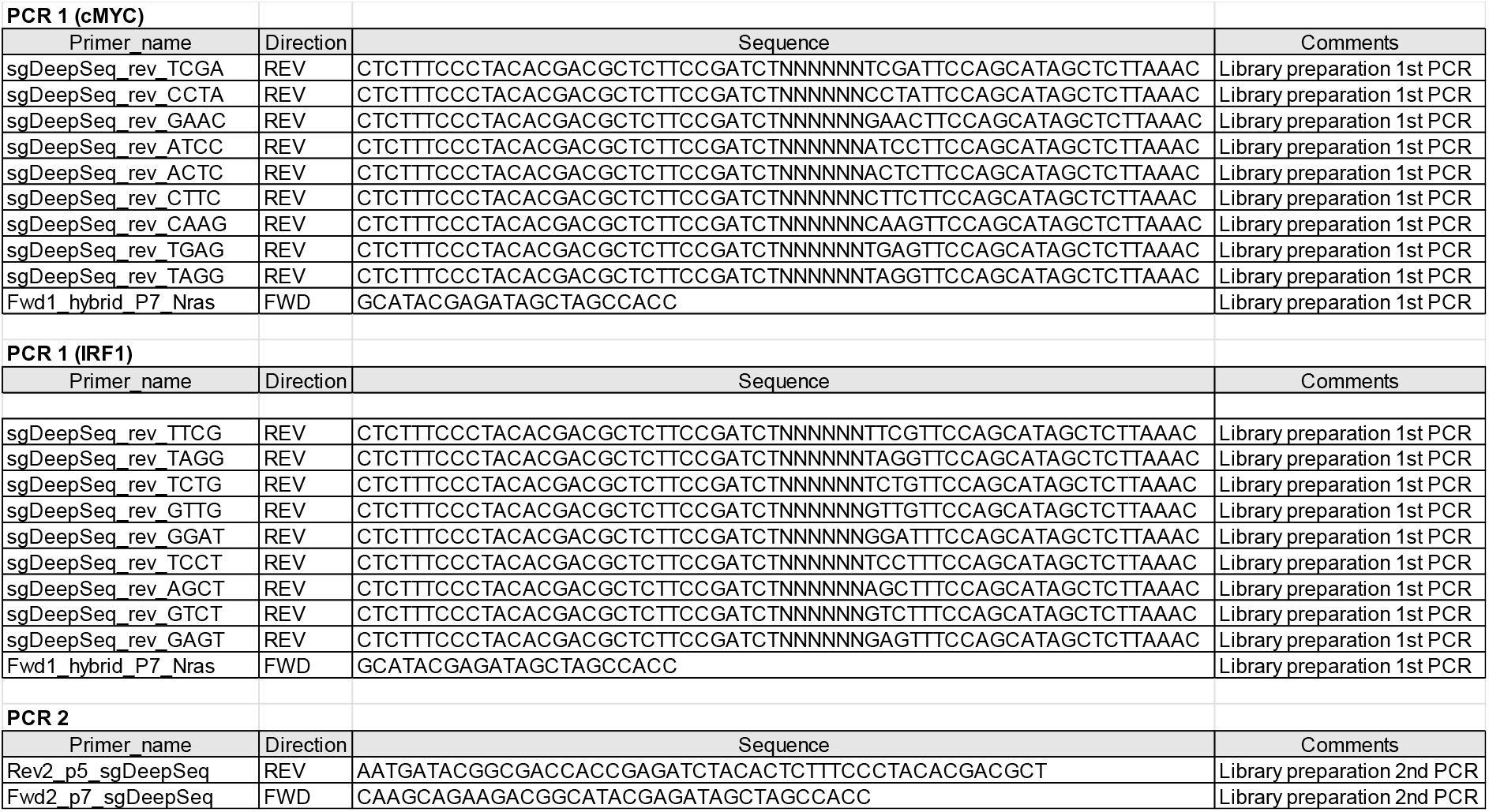

**Appendix figure 1.**
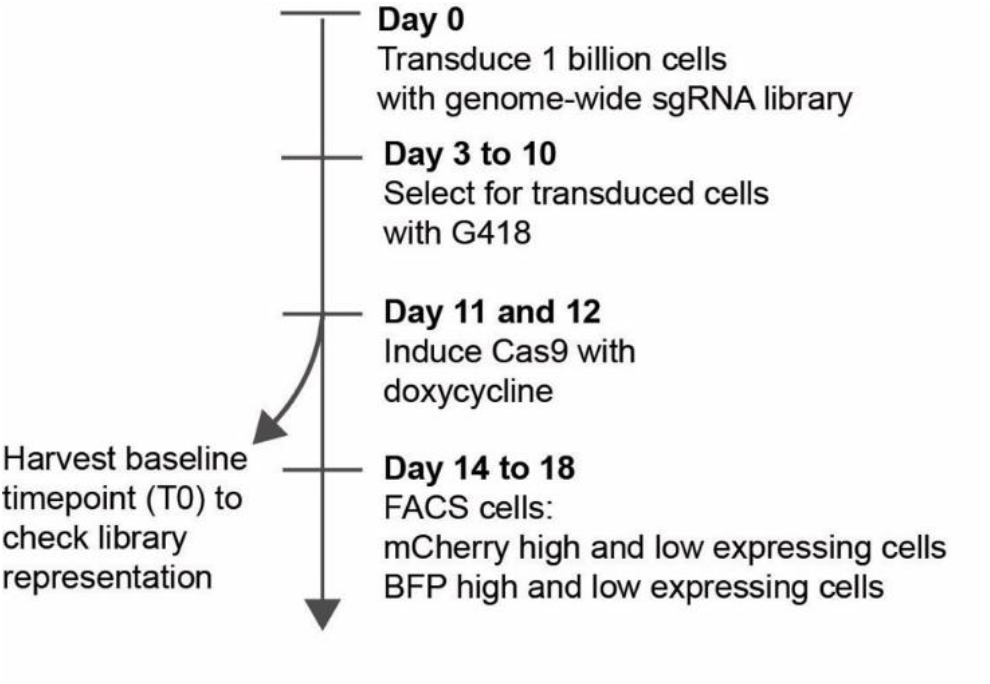
Time line and key steps in the genome-wide mCherry-cMYC and mCherry-IRF1 pooled forward genetic screens.

**Appendix figure 2.**
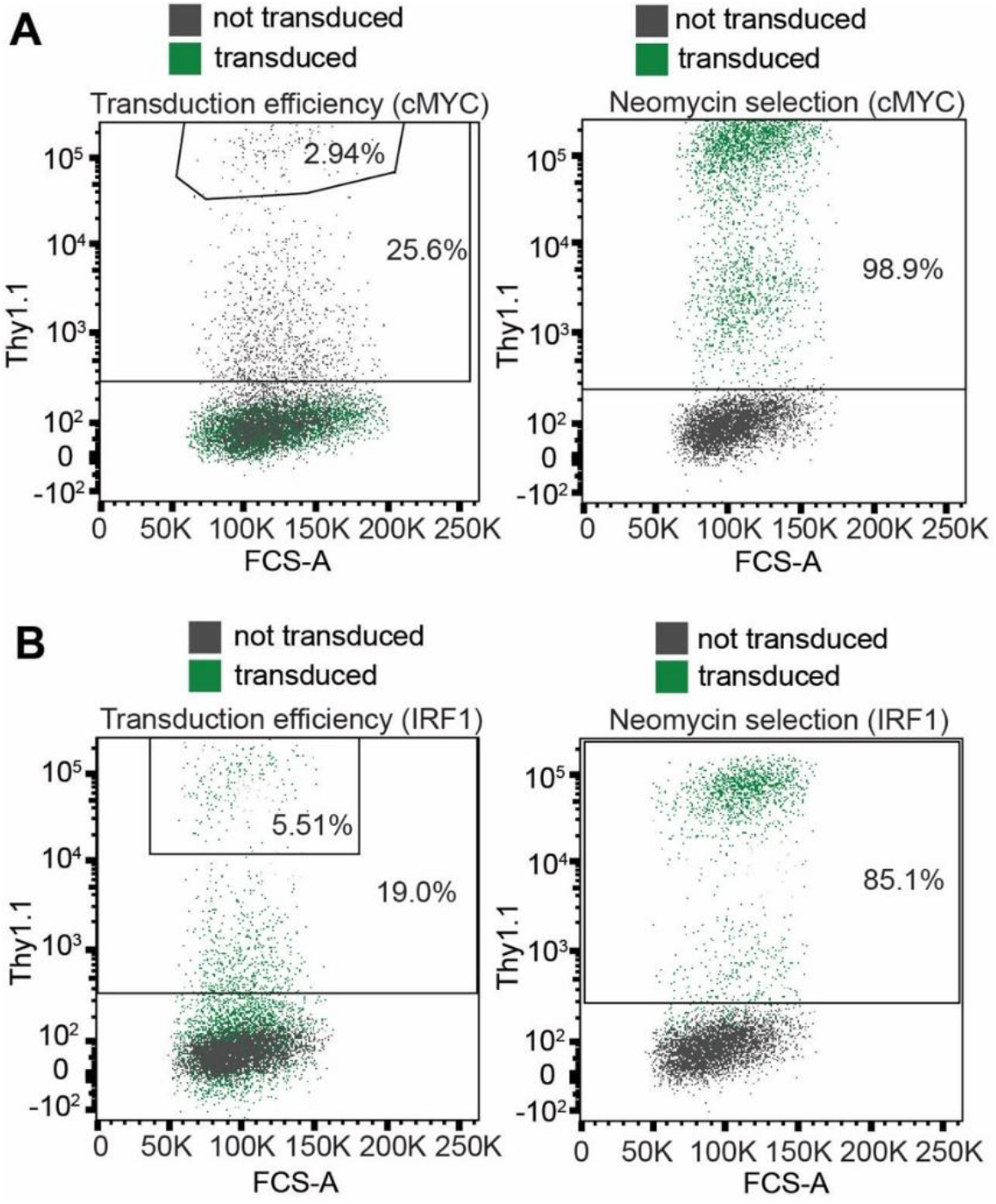
**A)** RKO-mCherry-cMYC-P2A-BFP cells, or **B)** RKO-mCherry-IRF1-P2A-BFP cells were transduced with a lentiviral genome-wide sgRNA library, harboring a Neomycin-resistance cassette. Transduction-efficiency before (left panels) and after (right panels) G418 selection was determined by Thy1.1 staining and flow cytometry.

**Appendix figure 3.**
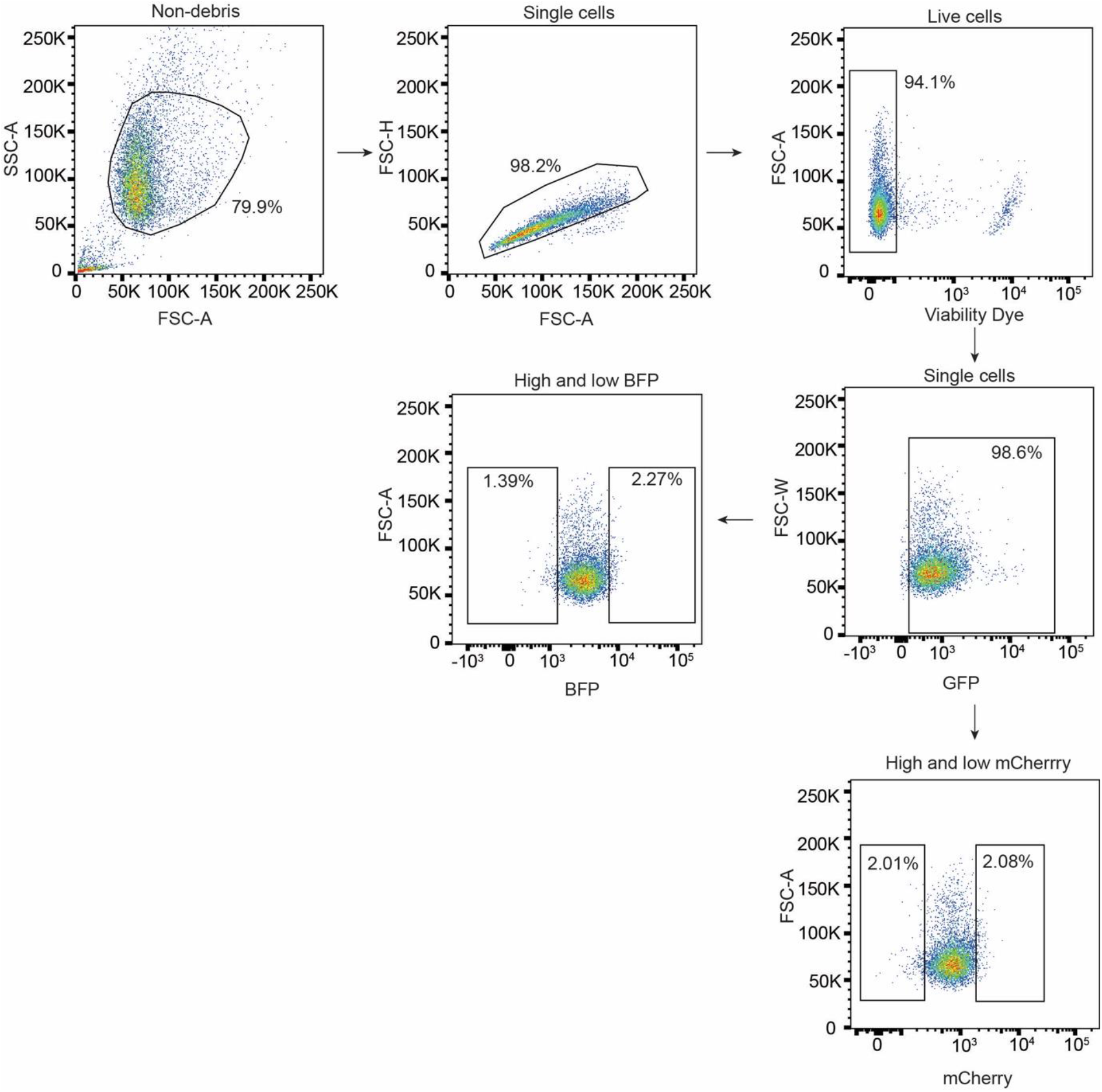
Gating strategy for the mCherry-cMYC screen.

**Appendix figure 4.**
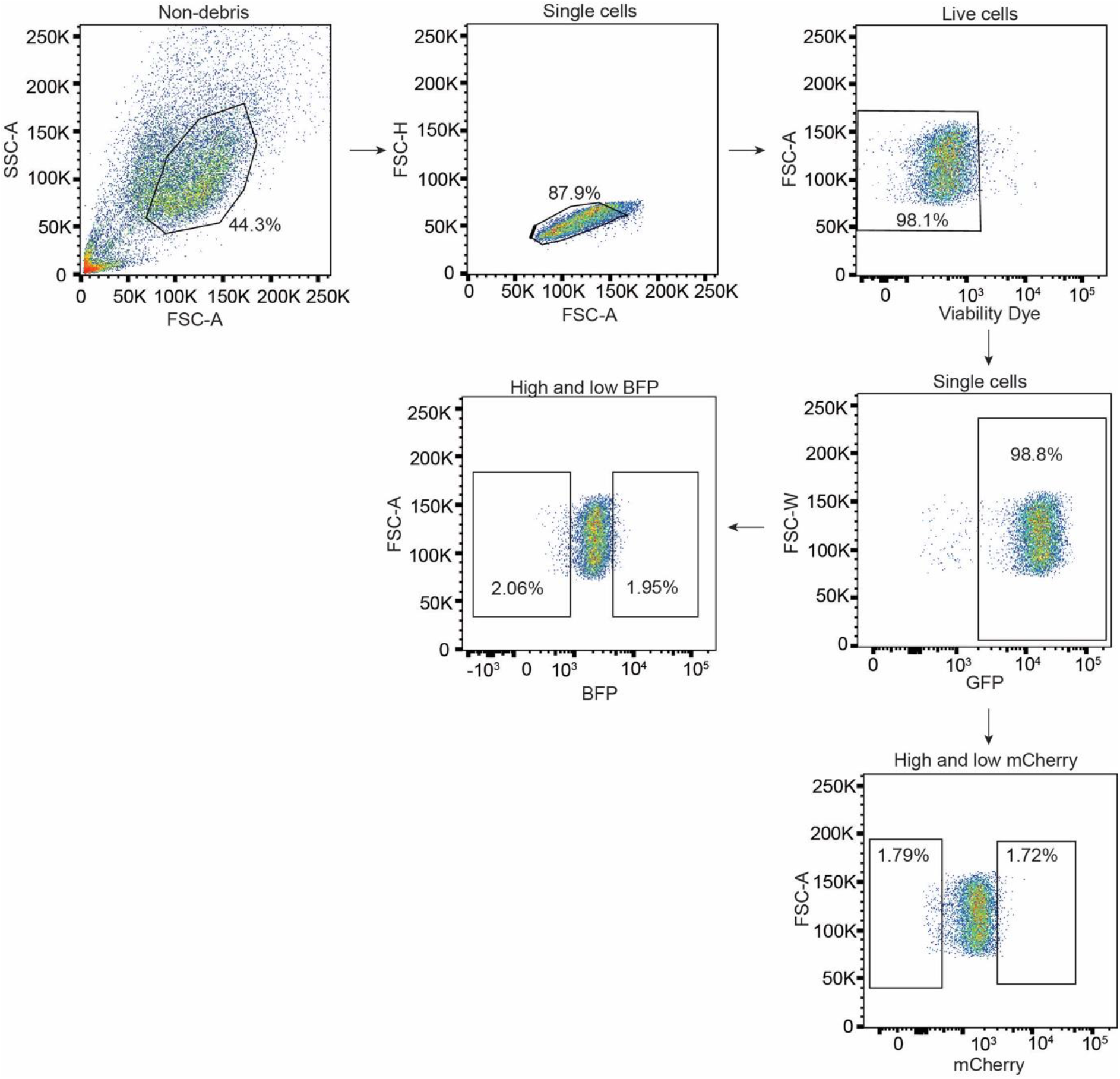
Gating strategy for the mCherry-IRF1 screen.

**Appendix figure 5.**
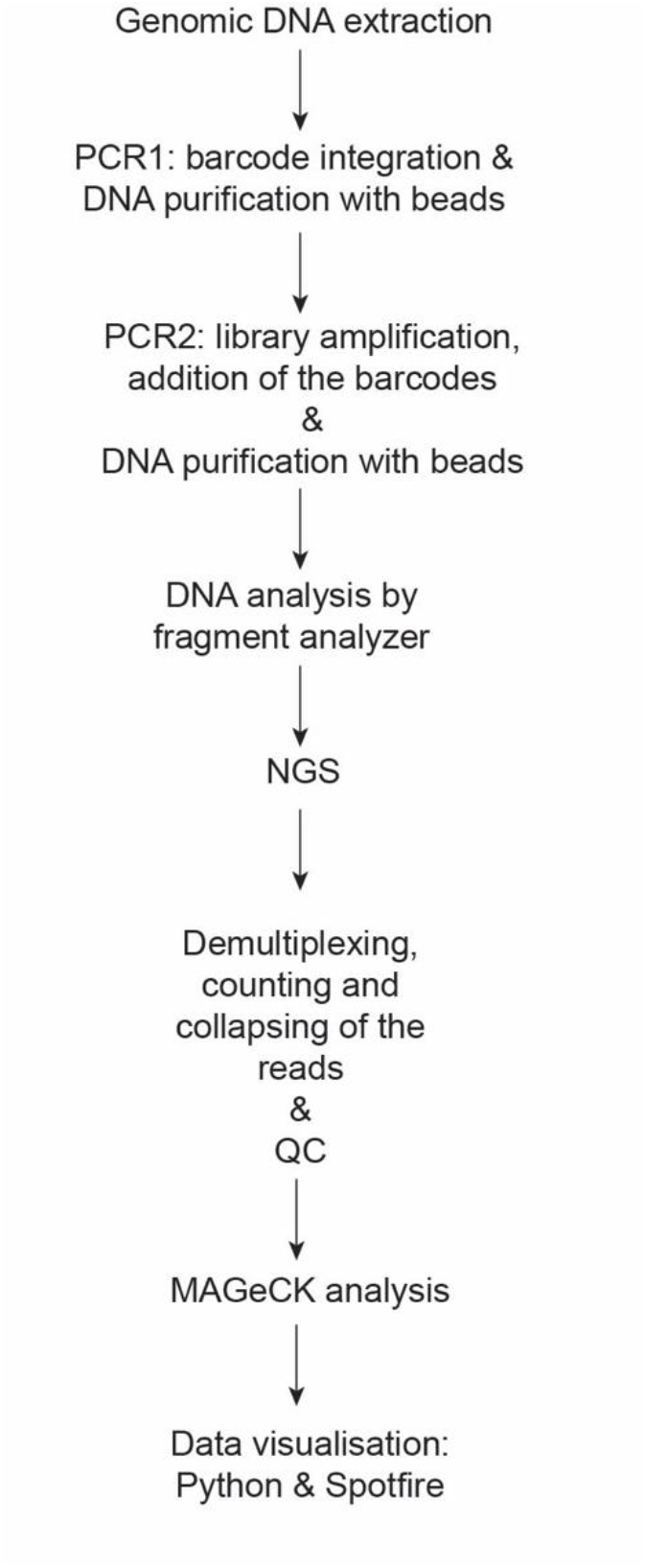
Flow chart for analysis of screen NGS data sets.

**Appendix figure 6.**
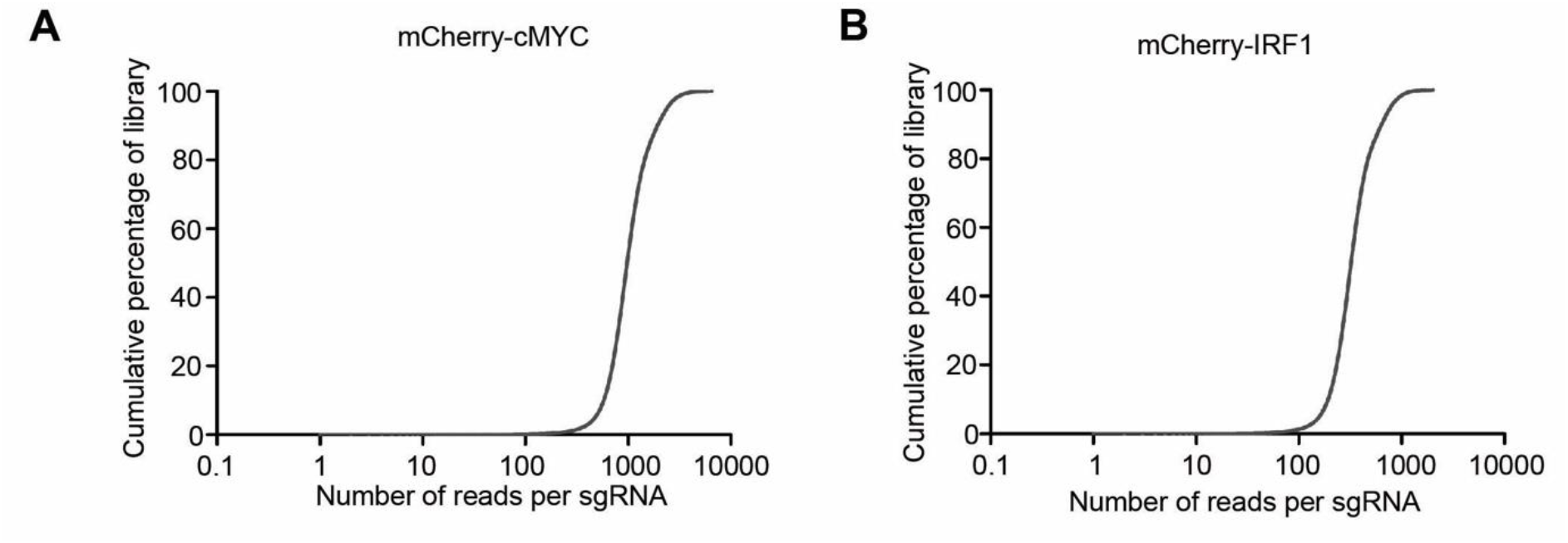
Cumulative percentage of library coverage per sgRNA coding sequence for the **A)** mCherry-cMYC screen and **B)** mCherry-IRF1 screen.

**Appendix figure 7.**
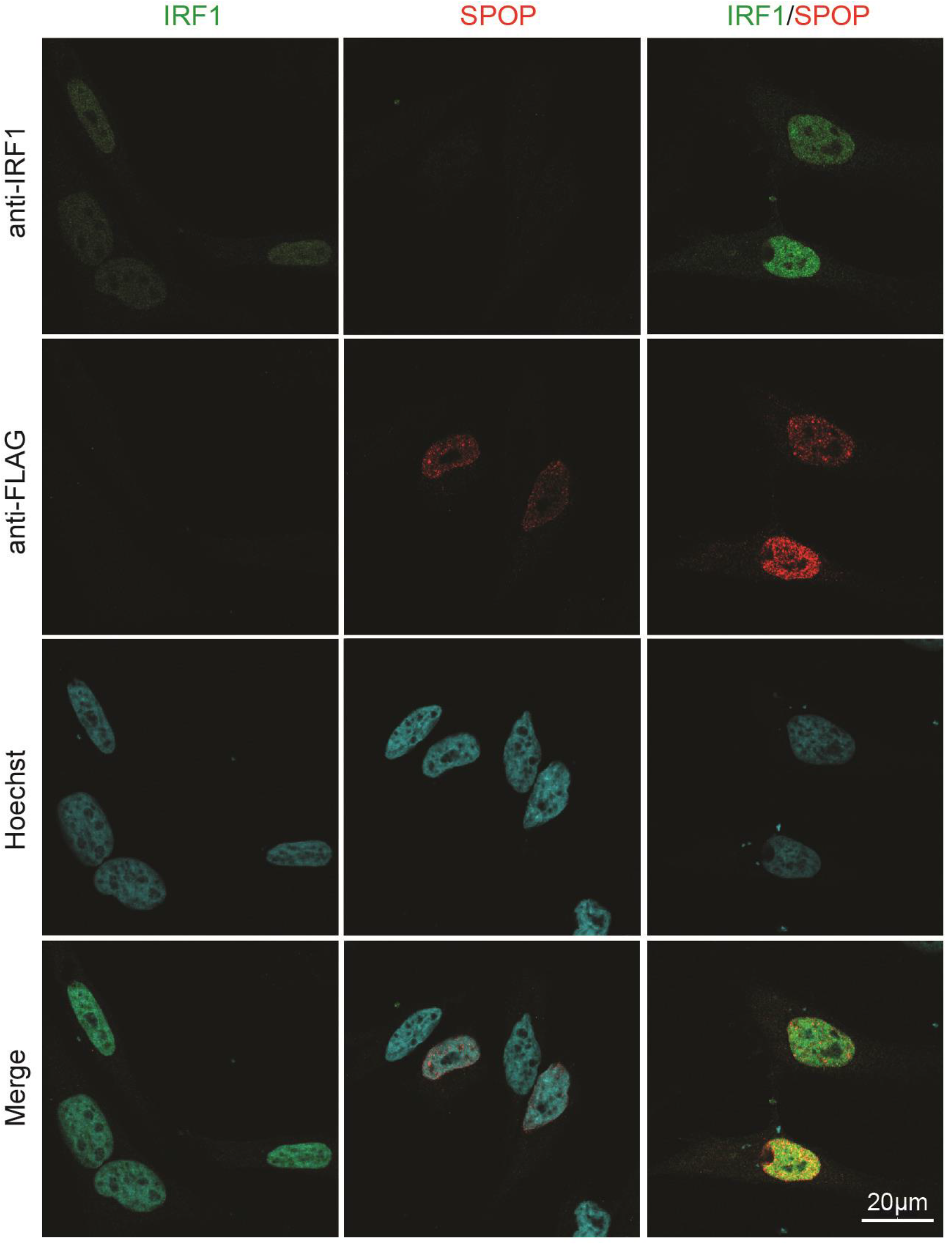
HeLa cells were transfected with expression plasmids for MYC-tagged IRF1 and FLAG-tagged SPOP. At 48 h. post-transfection, cells were fixed and localization analyzed by confocal immunofluorescence microscopy. Extended version of Fig. 4d, including single-transfected controls and single channels.

